# Thalamus sends information about arousal but not valence to the amygdala

**DOI:** 10.1101/2022.07.18.500471

**Authors:** Chris A Leppla, Laurel R Keyes, Gordon Glober, Gillian A Matthews, Kanha Batra, Maya Jay, Yu Feng, Hannah S Chen, Fergil Mills, Jeremy Delahanty, Jacob M Olson, Edward H Nieh, Praneeth Namburi, Craig Wildes, Romy Wichmann, Anna Beyeler, Eyal Y Kimchi, Kay M Tye

## Abstract

**Rationale:** The basolateral amygdala (BLA) and medial geniculate nucleus of the thalamus (MGN) have both been shown to be necessary for the formation of associative learning. While the role that the BLA plays in this process has long been emphasized, the MGN has been less well-studied and surrounded by debate regarding whether the relay of sensory information is active or passive.

**Objectives:** We seek to understand the role the MGN has within the thalamoamgydala circuit in the formation of associative learning.

**Methods:** Here, we use optogenetics to dissect the MGN-BLA circuit and explore the specific subpopulations for evidence of learning and synthesis of information that could impact downstream BLA encoding. We employ various machine learning techniques to investigate function within neural subpopulations. We introduce a novel method to investigate tonic changes across trial-by-trial structure, which offers an alternative approach to traditional trial-averaging techniques.

**Results:** We find that the MGN appears to encode arousal but not valence, unlike the BLA which encodes for both. We find that the MGN and the BLA appear to react differently to expected and unexpected outcomes; the BLA biased responses toward reward prediction error and the MGN focused on anticipated punishment. We uncover evidence of tonic changes by visualizing changes across trials during inter-trial intervals (baseline epochs) for a subset of cells.

**Conclusion:** We conclude that the MGN-BLA projector population acts as both filter and transferer of information by relaying information about the salience of cues to the amygdala, but these signals are not valence-specified.

**Preface:** In tribute to Nadia Chaudhri, and her discoveries regarding how contexts can modulate the representation of cues in the amygdala (Sciascia et al., 2015), and other mesolimbic circuit components (Chaudhri et al., 2009, 2008; Valyear et al., 2020), here we explore what information processing precedes entry into the basolateral amygdala (BLA) from the thalamus and develop visualization approaches for exploring across-trial temporal dynamics in addition to within-trial population activity.

## Introduction

Many similarities exist between the medial geniculate nucleus of the thalamus (MGN) and the basolateral amygdala (BLA) in how each region encodes information (Weinberger, 2011). Not only are both regions necessary for the formation of cued fear learning, but they both exhibit plasticity following learning, and encode information of multiple sensory modalities (Edeline et al., 1990; Wepsic, 1966). Numerous studies have focused on the BLA for associative learning (Nishijo et al., 1988; Weinberger, 2011), and for fear and reward learning (McKernan and Shinnick-Gallagher, 1997; Namburi et al., 2015; Rogan et al., 1997).

However, few studies have explicitly tested associative learning in the MGN (Bordi and LeDoux, 1994; Cruikshank et al., 1992; Edeline et al., 1990), due to the long-held assumption that the MGN only passively contributes to learning by relaying crude tone and somatosensory information (LeDoux, 2000; Orsini and Maren, 2012). However, there is evidence that the MGN likely plays a dynamic role in the formation of associative learning (Edeline et al., 1990; Parsons et al., 2006), fear conditioning (Ferrara, 2015; Nabavi et al., 2014) and fear memories (Antunes and Moita, 2010; Han et al., 2008). It has been argued that the MGN is the “root” of the auditory associative learning circuit, playing an active role and complimenting the BLA in the formation of cued fear memories (Weinberger, 2011).

The current work explores mechanisms underlying the formation of associative learning for both reward and punishment-associated outcomes by evaluating the similarities and differences between the MGN and the BLA neural population responses. We use a variety of approaches, such as hierarchical clustering and neural trajectories, to identify categorical differences in how these regions process discrimination learning. In order to do so, we dissect the neural circuit connectivity by exploring direct and indirect connections between the two regions. We use traditional experimenter-defined *a priori* methods to categorizing arousal and valence encoding patterns in each region. We further employ an algorithmic hierarchical clustering approach to uncover unbiased similarities and differences between how and what these regions encode. This work employs *in vivo* single-unit electrophysiology and circuit-specific optogenetics; the former allows us to characterize what information the MGN transmits to the BLA during Pavlovian learning and the latter provides a method to identify which cells in the MGN project to the BLA.

## Materials and Methods

### Experimental Design

#### Animal Care and Subject Details

Adult wild-type male C57BL/6J mice aged 8 weeks upon arrival were used (Jackson Laboratory; RRID: IMSR-JAX:000664). All experimental subjects were maintained in reverse light-cycle cubicles with *ad libitum* food and water until behavioral experiments began. Animals were housed four to one cage until electrode implantation, after which they were housed singly. All animal handling procedures were in accordance with those put forth by the National Institute of Health (NIH) and were approved by the Massachusetts Institute of Technology’s Institutional Animal Care and Use Committee (IACUC).

#### Stereotaxic surgery procedures

For all subjects, surgery was performed under aseptic conditions using a small animal stereotax (David Kopf Instruments, Tujunga, CA, USA) and body temperature was maintained using a heating pad. Anesthesia was induced using a 5% mixture of isoflurane and oxygen. Following induction this mixture was reduced to 2-2.5% and was maintained throughout the duration on the procedure (0.5L/min oxygen flow rate). Once subjects were adequately anesthetized, ophthalmic ointment was applied to the subject’s eyes, hair was removed from the incision site using hair clippers, and the area was scrubbed using 70% alcohol and betadine 3 times each in alternation, with an injection of 2% lidocaine under the skin for local anesthetic. All measurements for viral injections, electrode implants, or optrode implants were made with Bregma as the origin. Following surgery, subjects were placed in clean cages with water-softened mouse chow to recover. Cages were placed either under a heating lamp or on a heating pad to aid in recovery.

#### Viral Surgery

To record from the medial geniculate nucleus (MGN) neurons in a circuit-specific manner, a combination viral approach was employed. Following the general surgery procedures detailed above, an incision was made to provide access to the skull. After cleaning the skull, craniotomies were made above the basolateral amygdala (BLA) and the MGN. In order to selectively express channelrhodopsin-2 (ChR2) in BLA-projecting cells in the MGN, an anterogradely-traveling adeno-associated virus serotype 5 (AAV5) coding for ChR2 fused with an enhanced yellow fluorescent protein (eYFP) in a double floxed inverted open reading frame (DIO) under the control of elongated factor-1α promotor (250nL of AAV5-EF1α-DIO-ChR2-eYFP) was injected into the MGN (stereotaxic coordinates: −3.0mm anteroposterior, 1.75mm mediolateral, - 3.9mm dorsoventral). Concurrently, the retrogradely-travelling canine adenovirus-2 coding for cre-recombinase (400nL CAV2-cre) was injected into the BLA (stereotaxic coordinates: −1.6mm anteroposterior, 3.35mm mediolateral, −4.9mm dorsoventral). Injections were carried out using a 10uL micro syringe (36g beveled needle) driven at a rate of 0.05uL/min by a micro syringe pump and controller (Micro4; WPI, Sarasota, FL, USA). Viral material was allowed to penetrate tissue prior to needle extraction (10-15 minutes/injection site). Separate needles were used for each virus to avoid contamination and were flushed thoroughly with sterile water following the surgical procedure. AAV5 viral aliquots were obtained from the University of North Carolina Vector Core (Chapel Hill, NC). CAV2 viral aliquots were obtained from the Institut de Génétique Moléculaire de Montpellier, France.

#### Electrode surgery

Implantation of electrode and optrode arrays took place 2-3 months following viral surgery to allow for robust viral expression. Following the general surgery procedures detailed above, an incision was made to provide access to the skull, and the skull was cleaned and scraped to provide a stable surface for anchoring microelectrode arrays. A total of 8 craniotomies were made to accommodate implantation of three anchoring skull screws, one microelectrode wire bundle, one optrode (comprised of a wire bundle and optical fiber), a ground wire, and passive headbar anchoring. The headbar (1in x 1/8in x 1/8in aluminum square stock) was attached using cement (Adhesive Dental Cement C&B Metabond; Parkell, Edgewood, NY, USA). Cement dried prior to implantation of skull screws (15-20 minutes). Following skull screw implantation electrode and optrode arrays were lowered into the BLA (−1.6mm anteroposterior, 3.35mm mediodorsal, −4.9mm dorsoventral) and the MGN (−3.0mm anteroposterior, 1.75mm mediodorsal, −3.9mm dorsoventral), respectively. The multiarray ground wire was placed between the skull and brain surface in a craniotomy contralateral to electrode and optrode implantation sites. Cement was then applied to the screws, arrays, headbar and ground wire, and allowed to dry for a minimum of 45 minutes or until hard to the touch. A protective layer of dental cement was then applied to secure all array and headbar elements. A nylon suture was used to close the incision around the implanted device. Prior to removal from anesthesia subjects received subcutaneous injections of warm saline (1mL) and meloxicam (5mg/kg). Following surgery subjects were allowed to fully recover from anesthesia prior to being returned to the animal housing facility.

### *In Vivo* electrophysiological recording

#### Electrode construction

Multi-electrode arrays used for recording single-unit activity were custom designed and built by the experimenter (CL). Arrays consisted of two multi-channel single-wire probes and were constructed using Omnetics micro-connector plugs (Omnetics Connector Corp., Minneapolis, MN) as the main structural component. The optrode array used for recording in the MGN was connected directly to the plug using a plastic spacer. This array was comprised of 17 single wires (22uM stablohm wire, polyamide insulated; California Fine Wire, Grover Beach, CA) connected to a 300uM optical fiber using polyamide microtubing to align the wires. The second array was floating and constructed such that it could be moved independently from the connector and optrode array. This electrode array also consisted of 17 wires of the same material, and wires were aligned using a length of syringe needle cut to suit (2-3mm in length). The wires in both arrays were connected to the contacts of the Omnetics connector by hand, and wire insulation was stripped by hand using forceps. Each connection was then painted with conductive adhesive to ensure a good connection (Silver Print; MG Chemicals, Burlington ON). Both cyanoacrylate glue and epoxy were used to secure components. Both arrays were cut to length by hand using fine serrated tungsten scissors (Fine Science Tools, Foster City, CA), with optrode wire bundle extended beyond the tip of the optical fiber (500-800uM). All wires were gold plated to decrease impedance to 150-250MΩ using a 50/50 gold plating solution comprised of gold solution (Neuralynx, Bozeman, MT) and 1uM polyethylene glycol. Following gold plating, a low-impedance bare silver wire (California Fine Wire, Grover Beach, CA) was soldered to the final pin on the connector, and the connection was coated in dental cement.

#### In vivo single-unit electrophysiological recordings

Neural activity was recorded using an Open Ephys acquisition board (Siegle et al., 2017, J Neural Eng, PMID: 28169219) in conjunction with 32 channel Intan headstages (Intan Technologies, Los Angeles, CA). These data were recorded at a sampling rate of 30kHz, using a band pass filter to collect signals between 1 and 7000 Hz. Data processing was carried out using a combination of commercially available as well as custom written suite of software. Initial waveform detection and creation of files for cluster sorting was carried out using a custom algorithm in MATLAB (MATLAB; MathWorks, Natick, MA). Identified spikes were threshold using a 6 sigma criterion to reduce the probability of including spatially-attenuated multiunit signals and traces were aligned to the depolarization peak. Thresholded and aligned traces were then exported in the .*plx* format to be imported into Offline Sorter (Plexon Inc., Dallas TX), which was used for cluster sorting the spike data using principal component analysis (PCA). Behavioral events were recorded simultaneously with neural data using analog inputs to the Open Ephys acquisition board with the same sampling rate as neural data and were in some cases multiplexed to reduce the number of recording channels required.

#### Optogenetic stimulation

Optogenetic stimulation for phototagging experiments was carried out using a 473nm Diode-Pumped Solid-State (DPSS) laser (OEM Laser Systems, Draper, UT) connected to a patch cable using FC/PC connections (Doric, Quebec, Canada). Light power for phototagging was calibrated to ∼22mW at the beginning of each recording session using a Thorlabs light power meter designed for use with lasers (Thorlabs; Newton, NJ).

#### Photoidentification protocol

To identify BLA-projecting cells in the MGN as well as those cells in the BLA receiving either direct or indirect input from the MGN, we expressed ChR2 in the MGN→BLA projections using the circuit-specific viral approach detailed above. Following the conclusion of behavioral training, but within the same neural recording session, light was delivered into the MGN using a laser connected to the optical fiber implanted 500-800um above the recording site (473nm, 22mW). Light was delivered as a series of pulses of varying length and frequencies as follows: 1s pulses, 5ms 1Hz pulses, 5ms 10Hz pulses, and 5ms 20Hz pulses. A custom MATLAB script was used to identify phototagged cells in the MGN using the following criteria: signed rank test, p<0.001, and z-score > 3.5 within 10ms of light onset for 5ms 1Hz laser pulses. The 10ms photoresponse latency threshold was used since no cells exhibiting recurrent excitatory connections were observed during *ex vivo* whole cell patching experiments. In Network cells in the BLA were identified using the following criteria: Rank-Sum test, p<0.001, within 50ms of laser onset 5ms 1Hz pulses.

#### Electrolytic lesioning

Electrolytic lesions were used to verify the placements of microelectrode arrays. To carry this out, subjects were placed under anesthesia using the same procedure as that for stereotaxic surgery. Lesions were created by passing current through singles wires in the multi-array. In the MGN, lesions were made on the channels identified to have phototagged cells. In the BLA, between 4 and 6 channels were chosen randomly for lesioning. Lesions consisted of passing current through each channel for 10 seconds. Subjects were sacrificed 15-30 minutes following lesions to allow for gliosis to occur.

### *Ex Vivo* electrophysiological recording

#### Ex vivo determination of photoresponse latency (PRL) threshold

Viral surgeries for all subjects were carried out on a cage-by-cage basis, therefore one subject from each cage was selected for collecting PRL data, while the other 3 subjects were used in behavioral recording experiments. Patching experiments to determine PRL threshold were carried out concurrently with behavioral training to ensure that PRL data were collected from subjects that had undergone a viral incubation period of similar length to behavioral recording animals. This period was approximately 3 months for all subjects. For this, subjects were deeply anesthetized using sodium pentobarbital (200mg/kg; IP injection), and transcardially perfused using 20mL of ice-cold artificial cerebrospinal fluid (aCSF) modified as follows to preserve brain sections: (composition in mM) NaCl 87, KCl 2.5, NaH2PO4*H20 1.3, MgCl2*6H2O 7, NaHCO3 25, sucrose 75, ascorbate 5, CaCl2*2H2O 0.5 in ddH20 (osmolarity 322-326 mOsm, pH 7.20-7.30, saturated with carbogen gas 95 % oxygen, 5 % carbon dioxide). The subject’s head was removed, and the brain rapidly extracted. The brain was mounted on the vibratome stage using adhesive and placed in the slicing chamber submerged in partially frozen modified aCSF (Leica VT1000S, Leica Microsystems, Germany). 300um coronal sections were taken along the extent of the MGN (AP: ∼2.5-3.5mm posterior to bregma). Brain slices were placed in a recovery chamber at 32°C containing carbogen saturated aCSF (composition in mM: NaCl 126, KCl 2.5, NaH2PO4*H20 1.25, MgCl2*6H2O 1, NaHCO3 26, glucose 10, CaCl2*H2O 2.4 in ddH20; osmolarity 298-301 mOsm; pH 7.28-7.32) for at least 1 hour prior to initiation of recording. Following recovery, slices were transferred to the electrophysiological recording chamber.

#### Whole-cell patch-clamp Recordings

Slices were restrained in the recording chamber using a platinum and nylon “harp”, and were continuously perfused with oxygenated aCSF (2mL/min; 30-32°C). Slices were imaged through a 40x water-immersion objective using an IR-DIC enabled upright microscope (Scientifica, Uckfield, UK) equipped with a QImaging Retiga EXi camera (Q Imaging, Canada). Whole-cell patch-clamp recordings were taken using borosilicate glass capillaries (World Precision Instruments, Hertfordshire, UK), pulled on a P-97 puller (Sutter Instrument, CA, USA). When filled within internal solution, the resulting recording pipettes had resistance values of 4-6 MOhm (internal solution composition in mM: potassium gluconate 125, NaCl 10, HEPES 20, MgATP 3, and 0.1 % neurobiotin, in ddH20; osmolarity 287 mOsm; pH 7.3). Whole-cell patch-clamp recordings were made from ChR2-eYFP expressing MGN cells using pClamp 10.4 software (Molecular Devices, CA, USA). Analog signals were amplified using a Multiclamp 700B amplifier, filtered at 3 kHz, and digitized at 10 kHz with a Digidata 1550 (Molecular Devices, CA, USA). ChR2-eYFP expressing BLA-projecting cells in the MGN were located by illuminating the slice briefly with 470nM light (pE-100; CoolLed, Andover, UK). In order to confirm ChR2 expression in the recorded cell, a sustained 1s pulse of blue light was delivered while recording from the cell in voltage-clamp mode. If sustained inward current was observed, the cell was determined to express ChR2-eYFP. Cells were then exposed to 5ms pulses of blue light (10 pulses delivered at 1Hz, every 60s) while recording in current-clamp mode to observe light-evoked spiking behavior. Data were analyzed post hoc using Clampfit 1.4 software (Molecular Devices, Sunnyvale, CA). Latency to spike was determined for each cell by taking the mean time from pulse onset to spike (peak) for 30 responses to blue light pulses. Because no spiking response was observed in non-expressing neighboring cells, the PRL was set at 10ms (Figure 2h).

#### Histology and storage

Following the conclusion of recording experiments subjects were anesthetized using 90 mg/kg sodium pentobarbital and perfused transcardially with 20 mL of ice-cold lactated Ringer’s solution, followed by 20mL ice-cold paraformaldehyde (4%; PFA) in phosphate buffered saline (PBS). Brains were extracted and placed in 4% PFA for 24 hours. Tissue was then equilibrated in cryo-protectant solution (30% sucrose in PBS, w/v). 60um coronal slices were taken from the tissue using a sliding microtome (HM430; Thermo Fisher Scientific, Waltham, MA), and stored in PBS at 4°C.

#### Immunohistochemistry

Sectioned tissue was incubated in 1x PBS solution containing 3% Donkey serum in 0.03% Triton for 1 hour at room temperature. Sections were then washed using 1xPBS for 10 minutes, followed by incubation in a solution containing a DNA-specific fluorescent probe (DAPI; 4’,6-Diamidino-2-Phenylindole; 1:50,000 in PBS). Sections were then washed in 1x PBS 4 times, 10 minutes each. Sections were then mounted on slides for imaging using PVA-DABCO (Sigma-Aldrich; St. Louis, MO).

#### Confocal or epifluorescence imaging

Stained tissue slices were imaged using either a confocal laser scanning microscope (Olympus FV1000), or an epifluorescence microscope (Keyence BZ-X). Images were taken using a 10x objective lens. Following imaging, the images were evaluated to determine the location of viral expression as seen via fluorescence from the ChR2-eYFP. Recording sites were located using electrolytic lesion locations, which were identified using auto fluorescence produced by gliosis. In some cases, lesions or electrode extraction created holes in the tissues at the recording sites making probe location readily identifiable. Data from animals with wires outside the regions of interest were excluded from analysis. If lesions were correctly located in only one region, then only data from that region were used for that animal. Following imaging, slides were stored in slide boxes at room temperature in case additional imaging or validation was necessary.

### Pavlovian behavioral training paradigms

All subjects were placed on a food regulation schedule prior to the initiation of Pavlovian training (4g/day standard mouse chow, *ad libitum* access to water). For this Pavlovian paradigm, highly palatable Ensure nutrient drink was the rewarding unconditioned stimulus (US), and a mildly aversive airpuff to the subject’s face (∼25psi, 75ms) acted as the punishing US. On the first day of training subjects were placed in the experimental apparatus, using head fixation via a small aluminum headbar held in place by a modified stereotaxic holder. The initial training session was used to expose subjects to reward-related stimuli only while the infrared (IR) beam break used for lick detection was manually calibrated. In cases where subjects demonstrated reluctance in lick responding to presentations of the rewarding US, the experimenter would deliver US rewards directly into the subjects’ mouth. Following equipment calibration, subjects underwent Acquisition training before proceeding to Discrimination trials. Acquisition training consisted of interspersed conditioned stimuli (CS) and unconditioned stimuli (US) presentation of reward-association trials and free reward trials to encourage licking. Reward association trials consisted of presentation of a 4-second pure tone (8.5KHz and 14KHz, counterbalanced). This was followed by a variable (jitter) delay of 1-1.5 sec whereupon US delivery took place. CSs were separated by a 15±4s inter trial interval (ITI). The number of trials presented in each training session was dictated by the experimenter, who monitored the subject’s performance to infer motivation level. During the first acquisition session, if subjects exhibited a low rate of responding, the experimenter in some cases would also present free-reward trials during the session (these trials are not included in later analysis). Behavioral criteria determining subject performance are described below in the Statistical methods section. Subjects meeting acquisition criteria begin discrimination training the following day. Discrimination sessions were comprised of 75 reward trials and 35 punishment trials (4s, 65dB, 20s±4s ITI) and lasted 1-1.5 hours in length. The trial structure included jitter in US presentation timing following CS onset but before US delivery: in this case between 1 second and 1.5 seconds. If subjects did not perform correct anticipatory lick responses to >50% or anticipatory licking trials, learning was deemed unsuccessful.

### Statistical Methods

All computation was performed in MATLAB, unless explicitly stated otherwise.

#### Determining parameters for statistical comparison of ITI vs. anticipatory licking

To determine how many trials should be evaluated for statistical comparison between licking behavior during ITI periods and licking during anticipatory period, bootstrapping was performed on pilot behavioral data (n=6 mice). The number of trials necessary to find a significant result for the effect size of licking differences observed was calculated by finding the probability of identifying a significant result for 1 through 50 trial presentations. Significance was defined as having Student’s t-test with a p-value <0.05. Each condition was iterated 1000 times by randomly sampling the data. Under these conditions, we could detect a significant difference present in the data 93% of the time with 15 trials. By 20 trial presentations, the detection probability will be raised to 97%.

#### Statistical determination of learning behavior for experimental data

To measure behavioral performance during acquisition, the licking behavior during the anticipatory period (the time between CS onset and US delivery) was statistically compared to licking behavior during a period equal in length during the ITI period preceding each CS. If the Student’s t-test returned a significant result (p<0.01) and the raw number of licks during the anticipatory period was greater on average, the animal was determined to have successfully learned the CS-US association. This test was performed twice for each session using the first 20 and last 20 trials within the session. The number of trials (20) was determined using bootstrapping method described above. Animals meeting criteria moved into discrimination training on the next session. Discrimination sessions randomly presented CSs consisting of Ensure reward (CS-E) and Airpuff punishment (CS-A) trials. To measure behavioral performance during discrimination sessions, a statistical comparison of the difference scores for licking behavior during ITI and anticipatory periods; difference scores allowed licking behavior to be compared across re protocols as the baseline licking behavior varied between the two protocols. The difference scores were calculated by subtracting the raw number of licks during the anticipatory period from the raw number of licks during a period of equal length taken from the preceding ITI period for each trial. A Student’s t-test was used to compare the difference scores for CS-E and CS-A trials. This test was carried out three times for each session, using the first 20 trials, last 20 trials and a set of 20 randomly selected trials. If the test returned a result of p<0.01 and the raw number of licks for the rewarding CS-E trials was greater than the raw number of licks for the punishing CS-A trials on any of the three tests, the subject was determined to have successfully discriminated between the reward and punishment associated CSs.

#### Statistical determination of differences of task responsiveness and multimodal encoding within MGN and BLA

To determine cell responsiveness, a Wilcoxon Signed Rank-Sum test was applied using an experimental window of 0-100ms from stimulus onset and a baseline windows 3s from preceding ITI period; cell was deemed responsive to stimulus if resulting p<0.01. To determine whether cell was excited to or inhibited by cue, we computed the z-score response for baseline and experimental windows; cell was deemed *excited* if the average z-score response was greater than 0 (*inhibited* if less than 0). To determine the proportional differences between categories (CS, US, both, none), chi squared tests were applied; where data violated the assumptions of chi-squared test due to low *n* (*n*<5), we applied a Fisher’s Exact test. To identify the higher proportion of cells responding to experimenter defined categories, the Kruskal-Wallis test was used. If results were significant, Wilcoxon Rank-Sum post-hoc tests were performed pairwise, and Bonferroni corrected to identify which category was significantly higher.

#### Calculation of Neural Trajectories for time-averaged data

To explore the neuronal dynamics during reward and punishment stimuli, peri-stimulus time histograms (PSTHs) for each neuronal unit were computed at 50 ms bin width. PSTHs were separated according to stimulus (CS-E or CS-A) and cell type (MGN/BLA). We performed principal component analysis (PCA) to find the representative features for each cell type. Data were smoothed using a Gaussian-weighted moving average within a smoothing window of 25 previous bins. The first two PCs captured 77% variance in the MGN cells and 46 % in the BLA.

We also explored the dynamics depending on whether the behavior was correct or incorrect for each stimulus, yielding 4 conditions for each cell type. The four conditions are defined as hit (correct licking behavior during CS-E); miss (not licking during CS-E); false alarm (licking during CS-A); correct rejection (not licking during CS-A), where licking is measured during the anticipatory period, e.g., after the cue and prior to stimulus delivery. Dimensionality reduction was performed as above within a smoothing window consisting of 10 previous bins. These trajectories were projected on 3D space using the first three principal components, PC1, PC2, and PC3. To determine the mean length of trajectories, a leave-one-out approach was used, where the length was computed *k* times, where *k* is the number of subjects, then averaged across each group. To compute the distance between trajectories, the same leave-one-out approach was taken, this time computing the distance between any two trajectories and averaging across the groups.

#### Functional hierarchical clustering algorithm for phasic responses of trial-averaged data

Prior to clustering, data with NaN values were removed (n=1). The trial-averaged response (a.k.a. PSTH) was computed using 100 ms bin widths. Normalization consisted of z-scores, which were calculated using the mean and standard deviation during the baseline period (−2 sec to cue onset) individually for each neuron. Finally, the data were smoothed along the time dimension using a Gaussian-weighted moving average within a smoothing window of 10 prior bins. To compare reward and punishment trials, data were concatenated to calculate universal clusters, allowing for comparisons between and within CS-E and CS-A protocols. A hierarchical cluster tree was generated using Ward’s method, which uses inner squared distance to determine hierarchy using a Euclidean distance metric. A cutoff threshold value of 0.23 of the max value in the set was selected and used to determine clusters. Clusters containing less than 3 cells were discarded (for Ensure/Airpuff: 7 cells did not meet cluster criteria; for Hit/Miss/False Alarm/Correct Rejection: 4 cells did not meet cluster criteria). These clusters were then divided according to the different brain regions of interest, MGN non-Phototagged, MGN→BLA projections, Out-of-Network BLA, and In-Network BLA. Heatmaps plotted for each region represent the smoothed z-score input data; clusters for each region are color coded based on the original cluster tree. An identical approach was taken to preprocess hit/miss/false alarm/correct rejection data, concatenating each category appropriately by cell type; the same clustering parameters were applied.

#### Statistical analysis to measure clusters within each neural region

To determine if one cluster in the MGN contained significantly more cells than other clusters, we used a leave-one-out approach; we computed the number of cells in each cluster *k* times, each time leaving out one subject. We applied a Kruskal-Wallis ANOVA to the leave-one-out data to determine if there was a difference across all the clusters. If the ANOVA result was statistically significant, we applied post-hoc Wilcoxon Rank-Sum tests to the following subset of pairs: we selected the cluster with the highest mean number of cells and compared it to other clusters. To correct for multiple tests, we applied the Bonferroni correction by adjusting the significance level (alpha=0.05), dividing by the number of pairs being tested. This same approach was used for BLA data.

#### Functional hierarchical clustering algorithm for tonic changes to baseline periods

To identify tonic changes occurring during baseline or ITI periods, the following approach was taken: for each neuron and stimulus condition, three periods of interest were (1) an ITI baseline from −3 to −2 sec prior to cue onset, (2) a pre-cue region from −1 s prior to cue onset to cue onset and (3) a response window from cue onset to 1 sec past cue onset. For each time window, the number of spikes per trial were summed. This yielded (1) baseline spike count per trial (2) pre-cue spike count and (3) the response period spike for each neuron and stimulus condition. To normalize, the mean and standard deviations the baseline period were subtracted from the trial-averaged spike counts for both pre-cue and post-cue and for both CS-E and CS-A. Data were then smoothed over trials using a Gaussian-weighted moving average with a smoothing factor of 0.85. The resulting four z-scored datasets were concatenated prior to clustering. Hierarchical clustering was performed using Ward’s method on a Euclidean distance metric and the threshold set at 20% of the max value. Clusters containing less than 3 cells were discarded (for trial-based clusters: 5 cells did not meet cluster criteria). To compute whether a change to the firing rate occurred during the pre-cue period, data for each cluster were plotted as spike count by trial number. A regression line was fit to the spike count data. If the confidence intervals around the slope of the regression line did not contain 0, then a change to the baseline firing rate was determined to have occurred from the first trial to the last during the pre-cue period.

## RESULTS

### Experimental Paradigm

To achieve robust discriminative learning, we employed a two-stage Pavlovian paradigm that allowed mice to acquire a reward association (acquisition phase) before introducing punishing stimuli (discrimination phase) to ensure robust and consistent learning across animals. Discrimination sessions randomly presented conditioned stimuli (CS) consisting of Ensure reward (CS-E) and Airpuff punishment (CS-A) trials, pairing each with the appropriate tonal unconditioned stimuli (4s, 65dB, 20s±4s ITI). US delivery was randomly jittered during an interval of 1 to 1.5 s after CS onset to provide adequate time for subjects to exhibit reliable anticipatory licking throughout the first second of tone onset. The anticipatory licking period was defined as the period following CS onset and preceding US onset (yellow region in Figure 1a). The correct behavioral response to CS-E was anticipatory licking while that of CS-A was lick omission. Discrimination learning was considered successful when subjects demonstrated 90% correct responding to CS-E and 70% correct responding to CS-A, as well as success in the anticipatory licking control during a single session (Figure 1b). Lick behavior was recorded using an infrared (IR) beam break that was calibrated to each subject during Acquisition phase training (Figure 1c). Data (N=12) were bootstrapped to determine that 20 trial presentations were sufficient to correctly determine significance in behavioral data (Figure 1d).

**Fig. 1.**
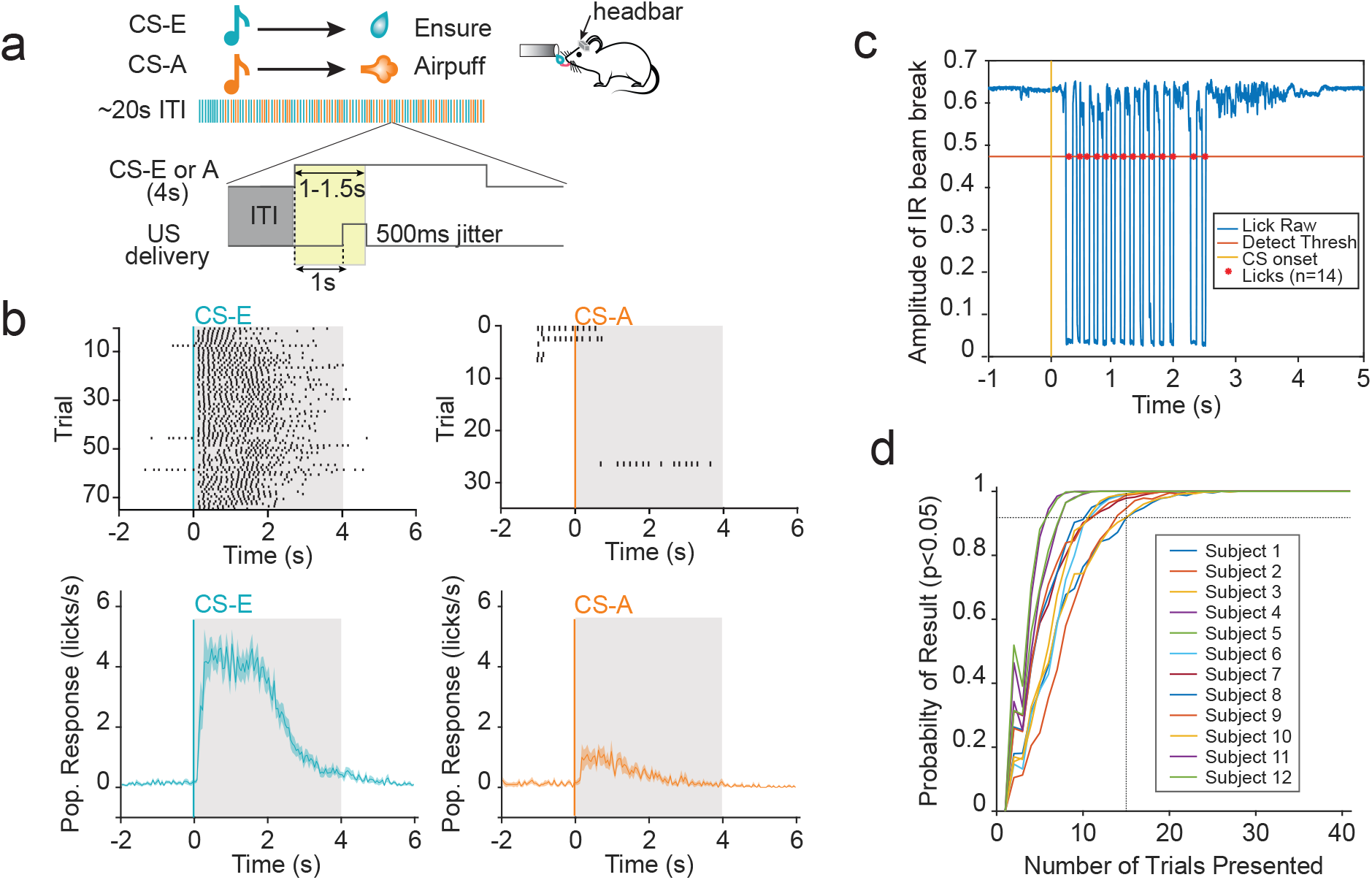
Behavioral lick response shows clear distinction between stimuli in experimental paradigm. (a) Schematic depiction of Pavlovian discrimination paradigm. Headfixed animals were trained to discriminate between two tones (conditioned stimulus, 8.5KHz and 14KHz); each tone was paired with a reward or punishment (Ensure or Airpuff, unconditioned stimulus) and counterbalanced across subjects. The series of colored bars shows pseudorandom trial order for reward (CS-E) and punishment (CS-A) experiments. The number of presentations per session are trial matched and comprised of 75 reward trials and 35 punishment trials (4s, 65dB tone, 20s±4s ITI). Detail shows schematic of time windows used for comparing ITI licking (gray) and anticipatory licking (yellow) for punishment experimental group. ITI window used was duration matched to jittered anticipatory period for each trial. (b) Representative licking response to presentations of reward-associated (top left) and punishment-associated CSs (top right) taken from one subject for each experiment. Gray shading represents 4s tone duration, black indicates licks for given time and trial. Mean normalized lick response with error for all subjects within each experimental group shown below. As expected with learned associations, licking behavior during reward trials (blue) is much higher than during punishment trials (orange). (c) Representative voltage trace (blue), from IR beam to detect licks. Individual lick events, indicated by red dots, show where the voltage drops below specified threshold. (d) Bootstrapping of behavioral data was used to determine the number of trials needed for sufficiently powered statistical analysis for the effect size of licking behavior during the anticipatory period vs. an ITI period of equal length. Subject data was taken from sessions of highest performance. Display shows the percentage out of 1000 samples that correctly detected a statistically significant difference (p<0.05) between the two licking periods. The resulting probability is above 90% using 15 trial presentations and above 98% using 20 trial presentations.

### Optogenetic stimulation allows identification of four distinct neural populations of interest: MGN non-Phototagged, MGN→BLA, Out-of-Network BLA, and In-Network BLA

To test the hypothesis that medial geniculate nucleus of the thalamus (MGN) neurons that project to basolateral amygdala (BLA) exhibit differences from MGN neurons during learning, we employed an optogenetic technique to identify specific subpopulations in the MGN and the BLA (Lima et al., 2009). We used a dual virus recombination approach to selectively express Channelrhodopsin-2 (ChR2) fused to a reporter, an enhanced yellow fluorescent protein (eYFP) in a single circuit: MGN neurons that project to the BLA (Figure 2a). By injecting an anterogradely-travelling virus carrying a cre-dependent ChR2 into the upstream brain region (MGN), and a retrogradely-travelling virus expressing cre-recombinase in the downstream region (BLA), only cells containing both viruses will express ChR2-eYFP, and thereby become light-sensitive and fluorescently labeled through cre-mediated specificity (Beyeler et al., 2016). Because the retrogradely travelling viruses are chosen for their propensity to enter neurons through axon terminals, viral recombination preferentially occurs in cells terminating in the downstream injection site (Fenno et al., 2011; Tye and Deisseroth, 2012). When ChR2 is activated by light, these non-specific cation channels open, causing an influx of ions into the cell body thereby causing a depolarizing current (Boyden et al., 2005). This depolarization can cause the cell to reach action potential threshold, causing the cell to spike (Rushton, 1927). These cells can then be identified as expressing ChR2-eYFP by recording the light-evoked spiking behavior, a powerful method known as phototagging. Furthermore, this technique can be combined with electrophysiological recording in multiple brain regions to identify other downstream neurons that are in the same connected network. Thus, by shining light into the upstream brain region and evoking responses there, we can also capture downstream responses; we then identify cells projecting to a downstream region by defining a short-latency response window to capture those cells firing in response to the light(Figure 2a, b).

**Fig. 2.**
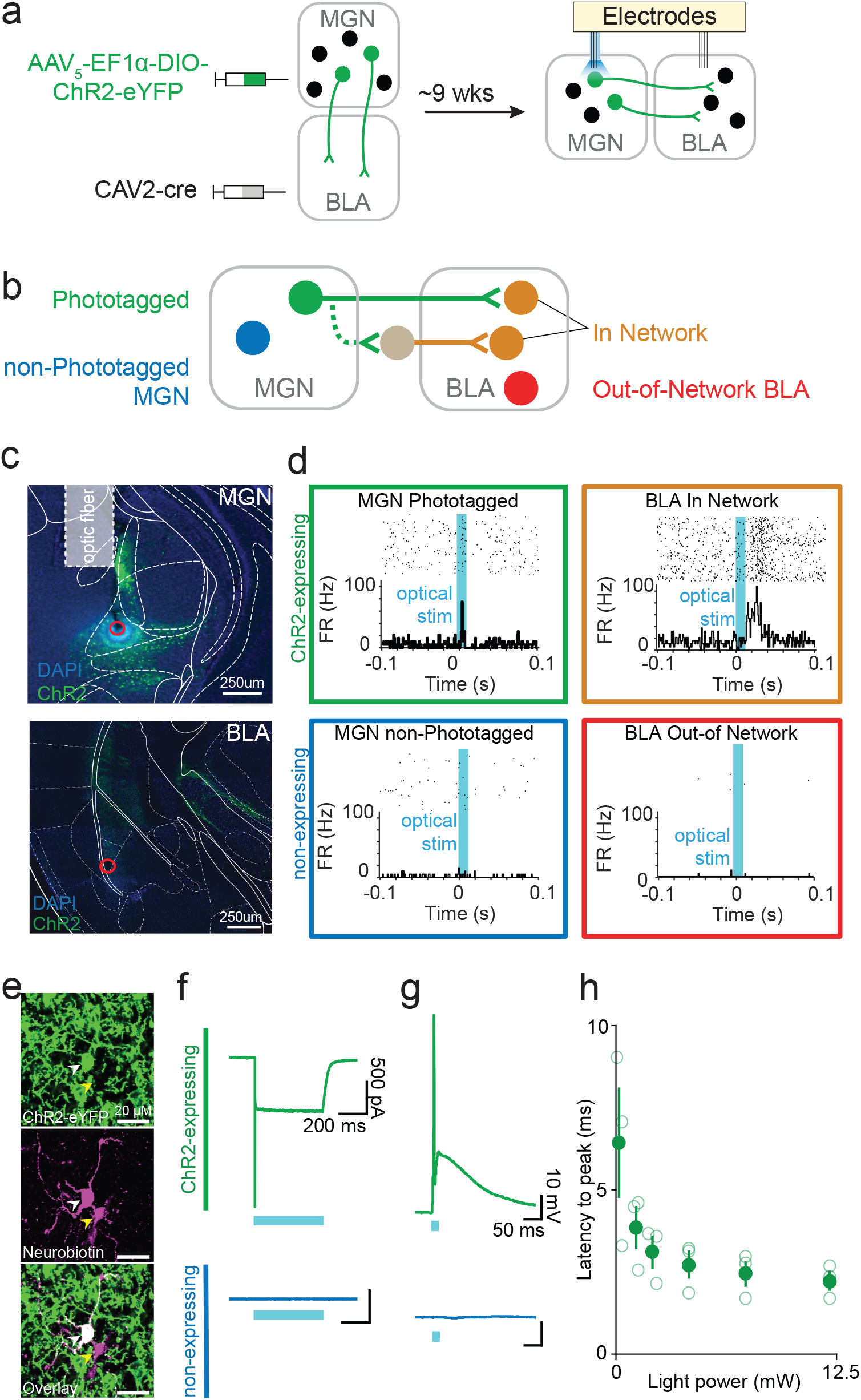
Phototagging viral approach and neural readout, *Ex vivo* evaluation of recurrent excitation and determination of photoresponse latency threshold. (a) Schematic depiction of combination viral approach for circuit specific expression of ChR2-eYFP in BLA-projecting cells in MGN. Injection volume: 250uL MGN, 400uL BLA. Schematic depiction of optrode and electrode placements within MGN and electrode-only placements within the BLA for collecting single-unit electrophysiology data from both regions simultaneously. (b) Schematic depiction of the four neural populations of interest: MGN→BLA projectors, unidentified MGN, in-network BLA, and unidentified BLA. Solid connections represent monosynaptic connections, whereas dashed connections represent synaptic connections including an unknown number of cells. (c) Representative images of combinatorial viral expression in MGN and BLA, as well as lesion sites in red. Illustrated optical fiber placement is approximate. (d) Representative photo-responses to blue light excitation (473 nm wavelength) for a Phototagged cell in MGN, an unidentified MGN cell, an In-Network BLA cell, and an Out-of-Network BLA cell. (e) Confocal images of neurobiotin-labelled neighboring ChR2-eYFP expressing and non-expressing cells in the MGN, following *ex vivo* electrophysiology. Overlay shows colocalization of ChR2-eYFP and neurobiotin in one cell (white marker) but not the neighboring cell (yellow marker). (f) Example traces, recorded in voltage-clamp mode, from ChR2-eYFP expressing and non-expressing cells during 1 second pulse of 473nm blue light. A ChR2-mediated sustained inward current is observed in the ChR2-expressing, but not in the non-expressing cell. A total of 13 cells were recorded from (n=10, non-expressing neighbors; n=3, ChR2-eYFP expressing cells). (g) Example current clamp recordings showing blue light-evoked ChR2-mediated spiking activity in the expressing cell but no response in the non-expressing cell. (h) Latency to action potential peak, from onset of light stimulation, as a function of light power for expressing cells. Each cell’s latency to peak was evaluated at 6 light power levels (0.2, 1.2, 2.1, 4.18, 7.38, and 12.12 mW/mm^2^).

Thus, the proposed circuit model represents the connectivity structure between the MGN and the BLA; it accounts for both the MGN neurons directly connected with the BLA identified via phototagging (MGN→BLA) as well as those polysynaptic connections for which only the BLA-projections (In-Network) are identifiable (Figure 2b). Representative images of lesion and viral expression showed that neurons achieved robust expression of ChR2 in the regions of interest (Figure 2c). *In vivo* neurons expressing ChR2 fire in response to blue light while neighboring neurons do not; thus, we can selectively stimulate cells in the MGN and evoke spikes in the downstream cells receiving input via the monosynaptic or polysynaptic connections (Allsop et al., 2018; Beyeler et al., 2018, 2016; Burgos-Robles et al., 2017; Nieh et al., 2015; Senn et al., 2014) (Figure 2d).

To validate our approach phototagging MGN→BLA projectors during single unit recordings, a photoresponse latency threshold was determined based on *ex vivo* whole cell patch-clamp recordings (N=3 mice; n=10 non-expressing neighbors; n=3 ChR2-eYFP expressing cells). Robust inward currents were observed in ChR2-expressing MGN cells during voltage-clamp recordings (Figure 2f), as well as short latency evoked spikes while recording in current-clamp mode (Figure 2g). Contrastingly, of the non-expressing neighboring cells recorded, no short latency light-evoked responses were observed (Figure 2e-g). This suggests the BLA-projecting subpopulation of the MGN does not have robust local recurrent connections. Having successfully validated the identification of the MGN→BLA projections via optogenetic stimulation, we recorded *in vivo* neural activity simultaneously in the MGN and the BLA using multiarray electrodes during behavioral tasks (Supplemental Figure 1).

### MGN neurons predominantly code for arousal while BLA neurons code for both arousal and valence

To gain understanding about the order of processes implemented across the thalamoamgydala circuit, we examined neural responses of MGN and BLA neurons during the Pavlovian discrimination task described above. To compare across subjects, data were normalized using z-score transformation and plotted on two different timescales (Figure 3a, b). The responses to CS-E were notably different between brain regions. All MGN cells exhibited a strong excitatory response to the reward cue. In contrast, BLA cells showed an excitatory phasic response of short duration (∼150 ms) then appeared to cease firing for the duration of the cue (Figure 3a). Because we were primarily interested in the learned association, these analyses are limited to the first second of the cue, prior to delivery of the US. No significant differences were found between the average response during the first second of the MGN Phototagged and non-Phototagged cells to either stimulus (CS-E: Wilcoxon Rank-Sum Test, p=0.89; CS-A: Wilcoxon Rank-Sum Test, p=0.75). Similarly, among BLA neurons, no significant differences were found between the In-Network and Out-of-Network BLA neurons in response to either stimulus (CS-E: Wilcoxon Rank-Sum Test, p=0.54; CS-A: Wilcoxon Rank-Sum Test, p=0.30).

**Fig. 3.**
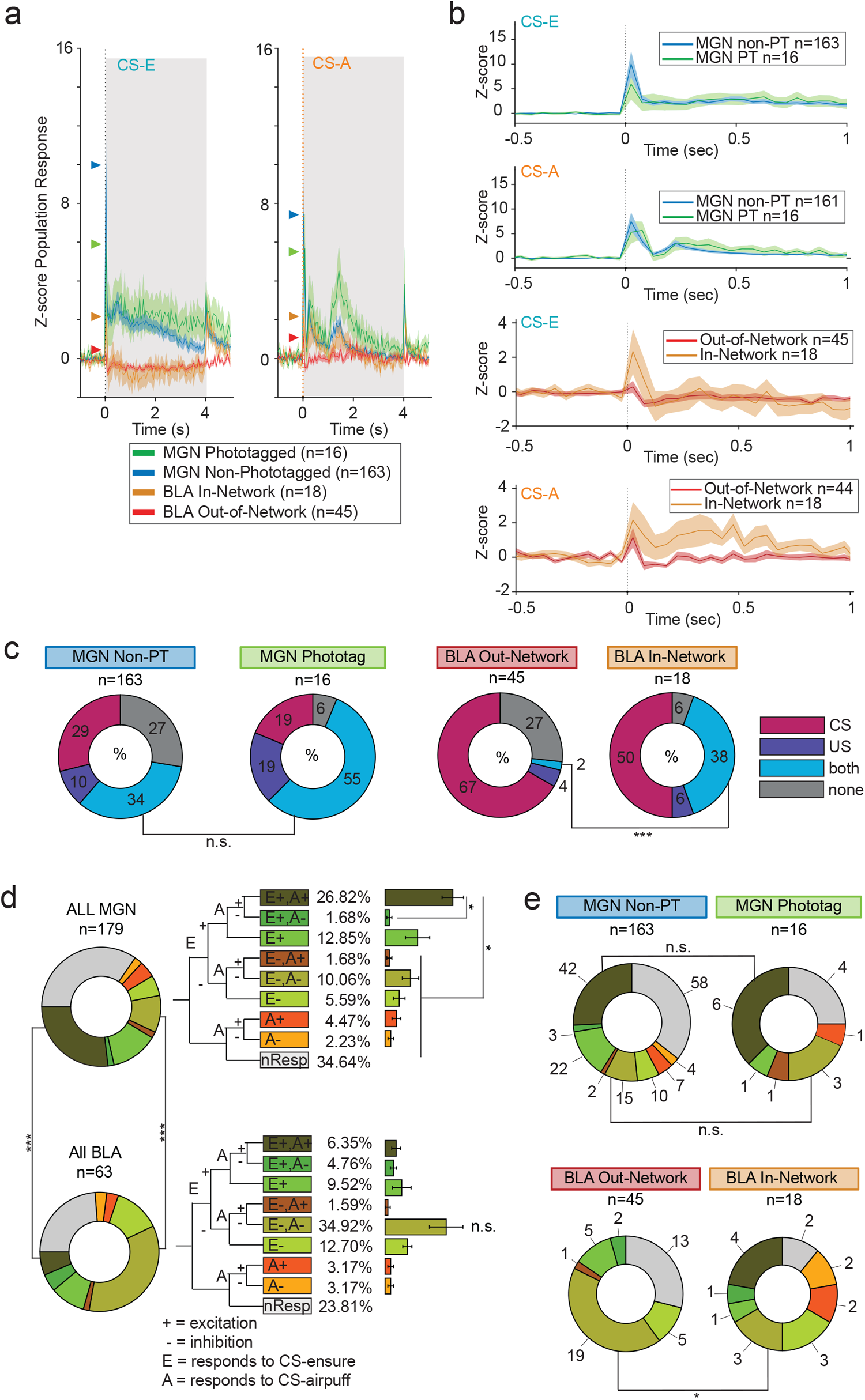
BLA-projecting MGN neurons are more likely to encode multimodal stimuli and be task-responsive than overall MGN population. (a) Z-score population responses to Ensure and Airpuff protocols over 6 s time window. Gray shaded area depicts 4 second CS tone, triangular markers depict peak response for each neural population. (b) Z-score population response over 1s time window to each protocol for the following population of neurons: MGN non-Phototagged, MGN→BLA, Out-of-Network BLA, and In-Network BLA. Shading shows SEM. (c) Breakdown by response category of each neural population during Discrimination task. The largest percentage of cells in MGN cells responded to both the CS and the US; there was not a significant difference in proportion between Phototagged and non-Phototagged populations responding to both CSs (X^2^ test, p=0.073). However, a significantly higher proportion of In-Network cells responded to both US and CS compared to Out-of-Network cells (Fisher’s Exact test, p=0.0004). Cells were defined as responding to a stimulus if p<0.01 for a Wilcox signed rank sum test, using experimental window = 0-100ms from stimulus onset and a baseline window = 3s taken from preceding ITI period. (d) Decision tree represents *a priori* experimenter-defined categorization of single-unit excitatory (+) and inhibitory (-) response to Ensure (E) reward and Airpuff (A) punishment-associated cue onsets. Population distributions represent categorical response for each cell type. Donut plots on left represent the breakdown of each cell population for tree categories defined to the right. Overall, the MGN contains a higher proportion of E+A+ responding cells than the BLA, while BLA contains a higher proportion of E-A-responding cells (Fisher Exact test: for E+A+ ***p=0.0036; E-A-****p=0.0001). Within the MGN, significantly more cells respond as E+A+ than other categories (Kruskal-Wallis test, ****p=6.1602e-07, E+A+ post-hoc comparisons to E+A-****p<0.0001, E+ n.s., E-A+ ****p<0.0001, E-A-***p< 0.003, E-****p< 0.0002, A+ ****p< 0.0001, A-****p< 0.0001). Within the BLA, the seemingly strong asymmetrical response to E-A-is not significantly different from other categories (Kruskal-Wallis test p < 0.03; E-A-post-hoc comparisons to other categories were not significantly different). A cell is determined as responding to reward or punishment-associated CS if p<0.01, signed-rank sum. A z-score threshold of zero was used to determine if the response was excitation or inhibition. (e) Donut plots represent the breakdown of each cell population for tree categories defined above; numbers represent raw cell counts. The largest number of cells across the MGN were excited by both stimuli. Similarly, most MGN→BLA projections across the population are excited by both stimuli and there is not a significant difference in proportion of E+A+ or E-A-Phototagged cells compared to non-Phototagged cells (E+A+: X^2^ Test, n.s.; E-A-: Fisher’s Exact Test, n.s.). In contrast, Out-of-Network BLA cells do not include the E+,A+ category at all (it is fully contained in the In-Network subpopulation) while the In-Network subpopulation contains proportionally fewer E-A-responding cells compared to the overall BLA (Fisher’s exact test, *p=0.048). Interestingly, the In-Network population also shows a greater variety of categories than the Phototagged subpopulation.

We found that although the MGN→BLA subpopulation had a trend towards higher levels of task responsiveness to both CS and US compared to the non-Phototagged MGN population, this difference was not significant at the 5% level (X^2^ Test, p<0.07; Figure 3c). However, significantly more BLA In-Network cells responded to both CS and US compared to the Out-of-Network population (Fisher’s Exact test, ***p < 0.005).

To further characterize the roles the MGN and BLA play in the formation of associative learning, we looked at the overall encoding strength in both regions with respect to both reward and punishment associated cues. To provide a categorical way of characterizing the entire population, neuronal response profiles were defined *a priori* as inhibitory or excitatory in response to reward and punishment CSs (Figure 3d). Of the cue-responsive neurons in the MGN, significantly more cells are excited to both CSs (E+,A+ category) than almost all other categories (Figure 3d inset: Kruskal-Wallis test, ****p=6.1602e-07, E+A+ post-hoc comparisons to E+A-****p<0.0001, E+ n.s., E-A+ ****p<0.0001, E-A-***p< 0.003, E-****p< 0.0002, A+ ****p< 0.0001, A-***p< 0.0001). However, no significant differences were found between categories in the BLA, after correcting for multiple comparisons (Figure 3d inset: Kruskal-Wallis test *p =0.027; E-A-post-hoc comparisons to E+A+, n.s., p=0.04; E+A-, n.s., p=0.03; E+, n.s., p=0.1; E-A+, n.s., p=0.02; E-, n.s., p=0.07; A+, n.s., p=0.02; A-, n.s., p=0.02). Moreover, there is a significantly higher excitatory response profile in the MGN to both cues compared to the BLA (Fisher’s Exact Test, E+A+ *** p=0.0036). In contrast, proportionally more BLA neurons have an inhibitory response to both reward and punishment than MGN neurons (Fisher’s Exact Test, E-A-****p=0.0001).

Notably, the highest proportion of Phototagged MGN cells across the population are contained in two response categories (E+,A+ and E-,A-) that do not discriminate between valence; they are excited to both CSs or inhibited to both. This suggests information relayed to the amygdala from auditory thalamus is largely salience or arousal rather than assigning valence. Taken together, these data are consistent with the notion that the thalamus is not assigning valence to the information, merely relaying its salience to the amygdala where valence is assigned.

To further dissect neural responses within our model of circuit connectivity, we explored the breakdown of responses across the subpopulations to each experimenter-defined *a-priori* category (Figure 3d, e). Despite the small sample size within categories, we observed the following points of interest: there is not a significant difference in the proportion across the non-Phototagged and Phototagged MGN neurons in either the E+,A+ nor in E-,A-categories (E+A+: X^2^ Test, n.s.; E-A-: Fisher Exact Test, n.s.). However, in the BLA, there is a significantly smaller proportion of In-Network cells in the E-A-category compared to the Out-of-Network population (Fisher’s exact test, *p=0.048). It is also interesting that the E+A+ category only appears in In-Network cells and is absent in the Out-of-Network population. Despite the low number of neurons, it appears that the In-Network BLA population is richly representing both salience and valence.

### Hierarchical clustering of neural responses to reward and punishment highlights greater variety of response profiles in MGN compared to BLA

To explore the dynamic neural response to stimuli across brain regions, we plotted neural trajectories, which show how the response evolves through time in reduced dimensional space (Figure 4a, b). The responses to each stimulus in the MGN population start off close together, then travel in separate paths following cue onset. While trajectories in the MGN have a roughly circular shape, CS-A returns to nearly the same starting point while CS-E does not complete the circle, ending farther away from the baseline starting point. This indicates a sustained difference from baseline firing in response to reward in the MGN. Here, the CS-A trajectory is longer than the CS-E trajectory indicative of a more dynamic response to the punishment cue (Wilcoxon Rank-Sum test, ***p=0.0029). The average distance between the two trajectories was measured over time; the distance, averaged across 1 s baseline (light purple), is significantly higher from the distance averaged 1 s after cue onset (dark purple) indicating that there are divergent dynamics in the MGN that occur following the cue (Figure 4a inset: Wilcoxon Rank-Sum test, ****p=5.3e-19).

**Fig. 4.**
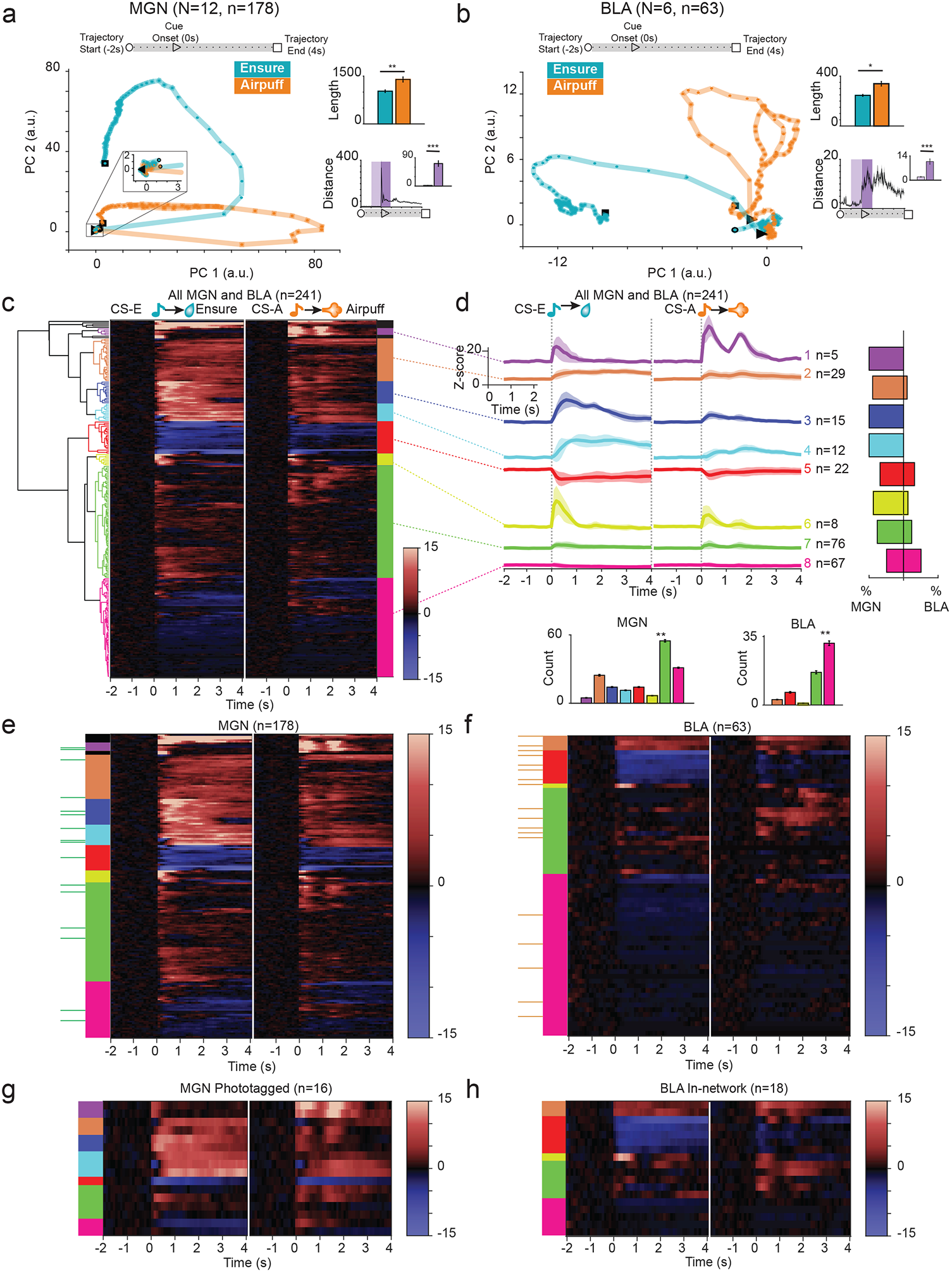
Clustering neural responses to stimuli exhibit more varied response pattern in MGN than BLA. (a, b) Neural trajectories along principal components for each condition are shown for MGN and BLA populations. The first three principal components explain 77% variance in MGN population. A pronounced jump following cue onset in MGN is clearly visible in the distance between the trajectories (inset Distance: light purple shows 1 s before cue, dark purple shows 1s following cue onset; Wilcoxon rank sum, p=5.3650e-19). Trajectories of BLA neurons (46% variance explained by first 3 PC) take a more varied path through the PC plane and a more dynamic response profile is seen for both stimuli compared to MGN. As in the MGN, there is a pronounced difference in the distance between trajectories following cue onset (Wilcoxon rank sum, p=1.6063e-18). In both regions, the CS-A trajectory is longer than the CS-E trajectory (inset Length: MGN: Wilcoxon rank sum test, p=0.0029; BLA: Wilcoxon rank sum test, p=0.0152). (c) Dendrogram shows relationship between 8 hierarchical clusters of neuronal response z-scores shown in heatmap to right. Heatmap shows response of all cells (n=241) to CS-E and CS-A protocols; tone onset at 0 s for each condition. Color bands to right of heatmap represent the 8 functional clusters output from hierarchical clustering (Ward’s method, Euclidean distance metric, cutoff height represents 23% of maximum data value). (d) Mean z-score response to CS-E and CS-A with error for all cells contained within each cluster; number of neurons contained within cluster is shown to right. Tone onset at 0 s for each stimulus. Scale shown on inset. Bar plot on right shows the breakdown of cell type as the total percentage of cells within each cluster that belong to MGN vs BLA. (e-g) Heatmaps arranged by cluster of single cell z-score responses to onset of CS reward and CS-punishment for each cell type: MGN non-Phototagged, Phototagged MGN→BLA, Out-of-Network BLA, In-Network BLA neurons. Cluster identity is shown using color coded key to left of each heatmap pair. Green ticks along far left side of MGN heatmap indicates phototagged cells and orange ticks along BLA show In-Network cells, which have been concatenated and shown in duplicate below each plot. Inset bar plots show the number of MGN or BLA cells within each cluster. Cluster #7 (green) is the most prevalent in the MGN (Kruskal-Wallis Test, ****p=4.3344e-17, Wilcoxon Rank-Sum post hoc tests comparing #7 cluster to: #1 (purple) ***p=0.00021; #2 (orange), p=0.00033; #3 (dark blue) **p=0.00031; #4 (light blue), **p=0.00029; #5 (red), **p=0.00031; #6 (yellow) **p=0.00028; #8 (pink) p=0.00032;). In the BLA, cluster #8 (pink) is the most prevalent response profile (Kruskal-Wallis Test, ****p=2.2658e-17, Wilcoxon Rank-Sum post hoc tests comparing cluster #8 (pink) to: #2 (orange) **p=0.00018; #5 (red), **p=0.00022; #6 (yellow), **p=0.00012; #7 (green), **p=0.00025). The MGN contains three clusters (#1-purple, #3-dark blue, and #4-light blue) indicating strong initial excitatory response and sustained excitatory response to stimulus that are not present in BLA. Interestingly, although the same clusters are present in MGN→BLA, they are not represented in the In-Network BLA. In a similar vein, the In-Network contains a cluster (#7-yellow) which is not seen MGN→BLA group.

In the BLA, again we find the total length of the punishment trajectory is significantly longer than reward, mirroring a more complex dynamic response in the BLA to punishment (Wilcoxon Rank-Sum test, **p=0.0152). The distance between trajectories increases following cue onset and there is a pronounced difference in the average distance over a 1 s baseline window compared to the 1s response window following cue (Figure 4b inset, Wilcoxon Rank-Sum, ****p=5.3650e-19).

We next performed hierarchical clustering to investigate neuronal responses without *a priori* determination of response type groups. This exploratory technique can reveal underlying structures that yield clues as to how these data are organized; the resulting clusters are evaluated for similarities and differences within and between experimental groups and brain regions. These resulting categories offer an alternative grouping than the experimenter determined categories in the previous section. Hierarchical clustering performed on 241 neurons aligned at cue onset over a 6s time interval for each protocol, yielded eight functionally distinct clusters (Figure 4c, d).

When broken down by cell type and subpopulation (Figure 4e-h), only five of eight clusters were common to both brain regions (#2 -orange, #5-red, #6-yellow, #7-green, #8-pink). The remaining three clusters (#1-purple, #3-blue, and #4-light blue) represent strong, excitatory responses; although these occur only occur only in the MGN, interestingly, all three are present in both Phototagged and non-Phototagged populations (Figure 4e, g). The cluster #7 (green) is the most prevalent in the MGN (Kruskal-Wallis Test, ****p=4.3344e-17, Wilcoxon Rank-Sum post hoc tests comparing #7 cluster to: #1 (purple) ***p=0.00021; #2 (orange), p=0.00033; #3 (dark blue) **p=0.00031; #4 (light blue), **p=0.00029; #5 (red), **p=0.00031; #6 (yellow) **p=0.00028; #8 (pink) p=0.00032;). In the BLA, cluster #8 (pink) is the most prevalent response profile (Kruskal-Wallis Test, ****p=2.2658e-17, Wilcoxon Rank-Sum post hoc tests comparing cluster #8 (pink) to: #2 (orange) **p=0.00018; #5 (red), **p=0.00022; #6 (yellow), **p=0.00012; #7 (green), **p=0.00025), showing a modest excitatory response to both CSs. From this analysis, we see a variety of responses specifically represented in the MGN Phototagged neurons that are providing information directly to the BLA; yet the In-Network BLA do not exhibit evidence of similar response profiles.

### MGN neurons as a population display more diversity in response profile compared with BLA neurons

To investigate the neural dynamics during specific signal detection responses, we examined the normalized neural activity in a reduced dimensional space using principal component analysis to visualize the neural trajectories (Figure 5a, b). Four conditions were defined using the licking behavioral responses to protocol stimuli, yielding the following combinations: Hit (licking during CS-E), Miss (not licking during CS-E), False Alarm (licking during CS-A), and Correct Rejection (not licking during CS-A). Neural trajectories within the MGN are well organized and smoothly varying according to condition, with false alarm, miss, and hit tracing similar patterns with slightly different lengths. The correct rejection population response travels along a different plane and is longer than the other three (Kruskal-Wallis test ****p<2e-07; Wilcoxon Rank-Sum post hoc tests: CR to Hit, ***p=4.7768e-04; CR to Miss, ***p=3.6585e-05; CR to FA, ***p=6.0058e-05). Correct Rejection is farthest from the Hit trajectory compared to the distance to Miss or to False Alarm (Kruskal-Wallis test, p=9.11e-05: Wilcoxon Rank-Sum post hoc tests Hit-to-CR compared to: Hit-to-Miss p=0.0058; Hit-to-FA, p=0.0086; CR-to-Miss, n.s.; CR-to-FA, n.s.; Miss-to-FA, p=0.0013). In the BLA, trajectory lengths are not significantly different when corrected for multiple comparisons (Kruskal-Wallis Test, *p=0.02; Wilcoxon Rank-Sum Tests, FA to Hit, p=0.0931, FA to CR, p=0.026, FA to Miss, p=0.026). The distance between False Alarm and Hit categories was greater than other pairings (Kruskal-wallis Test, p=1.11e-05; Wilcoxon Rank-Sum post hoc tests compared Hit-to-FA distances to: Hit-to-CR, n.s.; Hit-to-Miss, n.s.; CR-to-Miss, p=0.001; Miss-to-FA, p=0.0047). Qualitatively, the MGN appears to respond differently to a Correct Rejection compared with the other categories. However, the BLA appears to differentiate False Alarms from the other three conditions, given that False Alarm is the furthest away from the others. Although this False Alarm trajectory was not significantly longer in the BLA, clearly the dynamics are different.

**Fig. 5.**
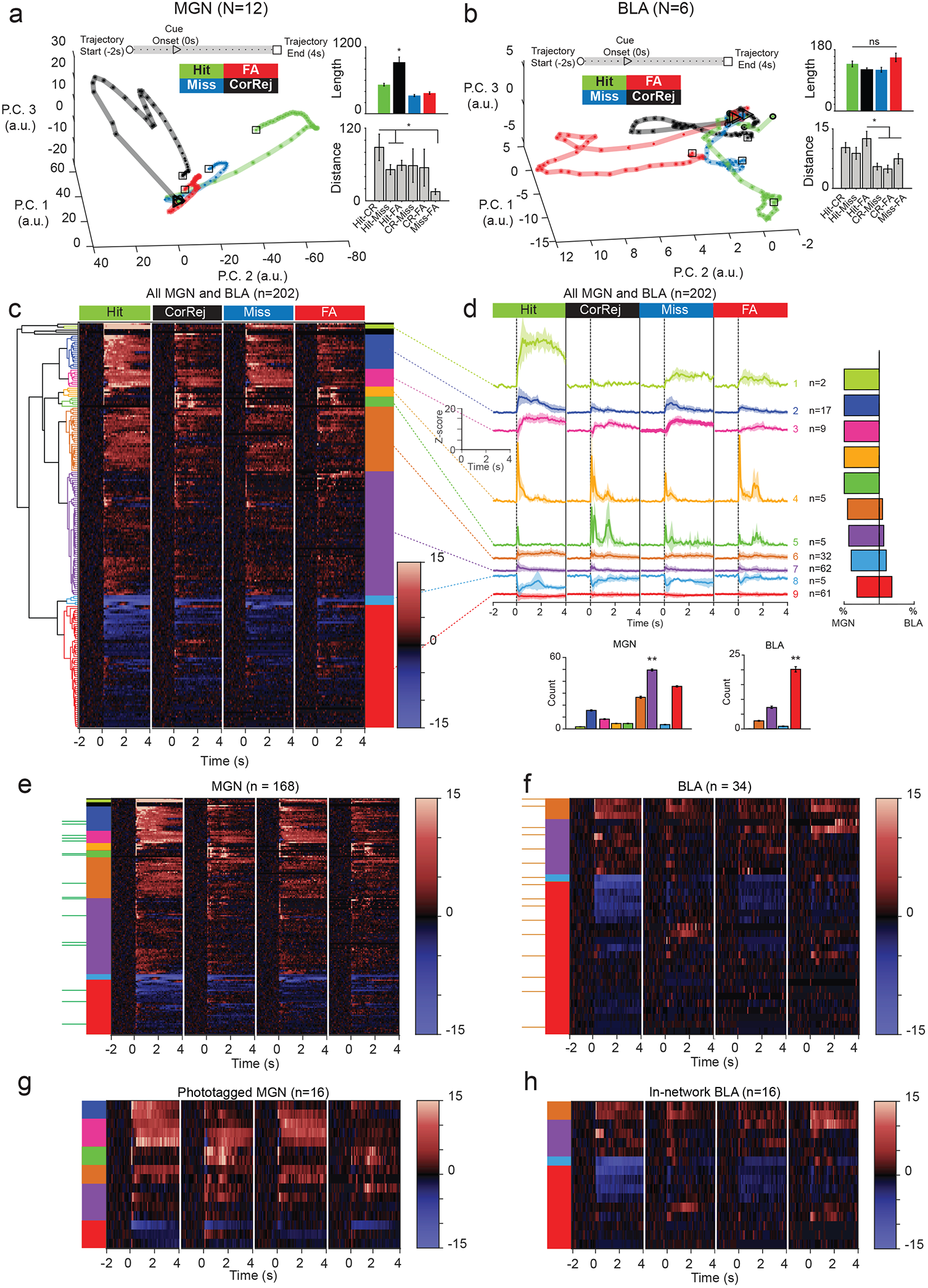
Clustering neural responses to Hit, Miss, False Alarm, and Correct Rejection behavioral categories have greater variety of response in MGN compared to BLA. (a) Neural trajectories along principal components for each condition are shown for MGN. Behavioral response categories are defined as Hit (licking when during reward), Miss (not licking during reward), False Alarm (licking during punishment), and Correct Rejection (not licking during punishment). In the MGN, Correct Rejection trajectory is the longest (Kruskal-Wallis test ****p<2e-07; Wilcoxon Rank Sum post hoc tests: CR to Hit, ***p=4.7768e-04; CR to Miss, ***p=3.6585e-05; CR to FA, ***p=6.0058e-05). This is reflected in the distances as well, with Correct rejection set apart from the other categories (Kruskal-Wallis test, ***p=9.11e-05: Wilcoxon Rank Sum post hoc tests Hit-to-CR compared to: Hit-to-Miss **p=0.0058; Hit-to-FA, **p=0.0086; CR-to-Miss, n.s.; CR-to-FA, n.s.; Miss-to-FA, **p=0.0013). (b) Neural response trajectories in the BLA using same categories defined in (a). Trajectory lengths are not significantly different when corrected for multiple comparisons (Kruskal-Wallis Test, *p=0.02; Wilcoxon Rank Sum Tests, FA to Hit, p=0.0931, FA to CR, p=0.026, FA to Miss, p=0.026). The distance between Hit-to-FA is longer than to CR-to-Miss and Miss-to-FA pairings (Kruskal-wallis Test, p=1.11e-05; Wilcoxon Rank Sum post hoc tests compared Hit-to-FA distances to: Hit-to-CR, n.s.; Hit-to-Miss, n.s.; CR-to-Miss, p=0.001; Miss-to-FA, p=0.0047). (c) Dendrogram shows relationship between 9 hierarchical clusters of neuronal response z-scores shown in heatmap to right. Heatmap shows response of 202 cells to each of 4 behavioral categories, tone onsets at 0 s for each condition. Color bands to right of heatmap represent the 9 functional clusters output from hierarchical clustering (Ward’s method, Euclidean distance metric, cutoff height represents 30% of maximum data value). (d) Mean z-score response of all cells grouped by cluster; cluster ID shown to right, number of neurons contained within cluster is shown to right of ID. Tone onset at 0 s for Hit, Miss, False Alarm, Correct Rejection categories. Bar plot on right shows the breakdown of cell type as the total percentage of cells within each cluster that belong to MGN vs BLA. (e-h) Heatmaps arranged by cluster of single cell z-score responses to onset of CS for each behavioral response category type for each population MGN, MGN→BLA, BLA, In-Network BLA neurons. Cluster identity is shown using color coded key to left of each heatmap pair. Green/Orange ticks along right side of MGN/BLA heatmaps indicate Phototagged/In-Network cells, which have been concatenated and shown in duplicate below each plot. Inset bar plots show the number of MGN or BLA cells within each cluster. All nine clusters are present in the MGN, the largest of which is cluster #7 (purple) (Kruskal-Wallis Test, p=7.29e-19; Wilcoxon Rank-Sum post hoc tests compare #7 (purple) to: #1 (light green) p=0.00033; #2 (dark blue) p=0.00032; #3 (hot pink), p=0.00027; #4 (yellow) p=0.00026; #5 (moss green) p=0.00021; #6 (orange) p=0.00033; #8 (light blue) p=0.00021; #9 (red) p=3.3e-5).). However, only four clusters are represented in the BLA, of which cluster #9 (red) is largest (Kruskal-Wallis Test p=3.76e-19; Wilcoxon Rank-Sum post hoc tests compare #9 (red) to: #6 (orange) p=1.7e-05; #7 (purple) p=2.02e-05; #8 (blue) p=1.7e-05). Although these same 4 clusters are represented in In-Network BLA subpopulation, there are only six of the 9 clusters represented in the MGN→BLA. As before, there is not a one-to-one match across categories between the MGN→BLA and In-Network BLA populations.

Performing hierarchical clustering on this set yields nine functional clusters (Figure 5c, d). The MGN overall represents all nine clusters (Figure 5e), of which cluster #7 (purple) is most prevalent (Kruskal-Wallis Test, p=7.29e-19; Wilcoxon Rank-Sum post hoc tests compare #7 (purple) to: #1 (light green) p=0.00033; #2 (dark blue) p=0.00032; #3 (hot pink), p=0.00027; #4 (yellow) p=0.00026; #5 (moss green) p=0.00021; #6 (orange) p=0.00033; #8 (light blue) p=0.00021; #9 (red) p=3.3e-5). However, only six of nine clusters are present in the Phototagged MGN subpopulation (Figure 5g). In the BLA only four clusters are present (Figure 5f), of which cluster #9 (red) is most prevalent (Kruskal-Wallis Test p=3.76e-19; Wilcoxon Rank-Sum post hoc tests compare #9 (red) to: #6 (orange) p=1.7e-05; #7 (purple) p=2.02e-05; #8 (blue) p=1.7e-05). In both In-Network and Out-of-Network populations, the same four clusters are present in both populations (Figure 5f, h). It is interesting to note that this pattern of cluster breakdown follows a similar pattern to the previous section (Figure 4g, h). There, we observed more variety in response profiles in the MGN with some overlap between MGN→BLA clusters. However, there was far less representation across clusters in both In-Network and Out-of-Network BLA.

### Between-trial hierarchical clustering reveals that a subset of neurons exhibit tonic changes during learning

Switching between tonic and burst firing patterns allows the thalamus to relay information to the cortex for efficient processing (Murray Sherman, 2001). We wondered whether there were differences in tonic and burst firing patterns in the thalamoamygdala circuit as well. We explored the inter-trial interval (ITI) for tonic changes during learning in concert with the strong phasic response following cue onset. To explore trends in the population response across the two brain regions, we examined the spike activity of all simultaneously recorded neurons. We computed the average z-score response for each raster, maintaining the trial structure by aligning each trial to the CS. The MGN population has robust phasic responses to each CS-E resulting in higher magnitude overall during the 1s response window (Figure 6a). In contrast, the BLA population exhibits mildly inhibitory responses to the reward cue (Figure 6b). In response to the punishment cue, the MGN exhibits a strong excitatory response immediately following the cue whereas the BLA exhibits an initial inhibitory response (Figure 6c, d). There is no evidence of tonic changes across the population for either stimulus, e.g., as might be seen by a steady change across baseline or response windows.

**Fig. 6.**
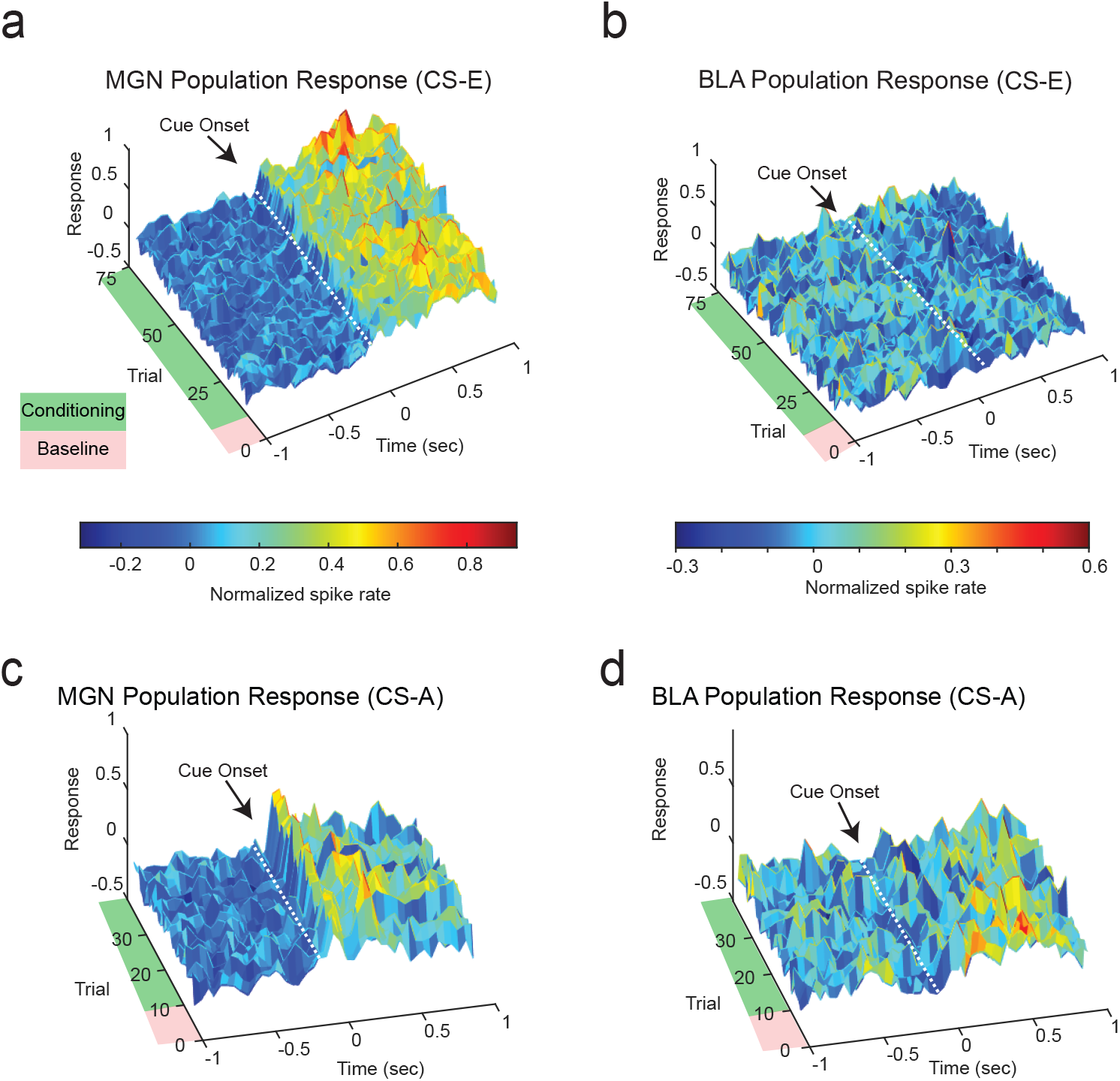
Population Dynamics Show Strong Responses to Audio Cue and No Visible Tonic Changes during Pre-Stimulus (Baseline) Periods. (a, b) Temporal dynamics of neuronal MGN and BLA population response to reward-predictive cue during discrimination trials. Spike activity of all simultaneously recorded MGN (BLA) neurons; 50 ms bins. Consistent with the small number of task-responsive neurons that change with conditioning (Supplementary Figure 2 D), conditioning does not produce a visible change in firing rate for either population. Note that conditioning also does not produce a visible change to the tonic firing rate during the pre-stimulus period. (c, d) Temporal dynamics of neuronal MGN and BLA population response to punishment-predictive cue during discrimination trials. Spike activity of all simultaneously recorded MGN (BLA) neurons; 50 ms bins.

During associative learning tasks, trial-to-trial changes during the response window are expected and are often taken as evidence of learning. However, the pre-cue (baseline) periods that are typically not expected to exhibit trial-by-trial changes; we expect the firing to be relatively static during pre-cue periods across trials. To test for subtle changes to the tonic firing rate during the pre-CS periods, we denoted a baseline period as 1s period before cue onset and an experimental response period of 1s response following cue onset for every trial. Rather than averaging across trials, we instead summed the number of spikes across trials within each window, yielding the spike count per trial across each time interval (Figure 7a). This allowed us to compare the number of spikes for each trial before and after the cue to identify trends. We note that this approach is unlike the one taken in Figures 4 and 5, where clustering was done using standard peri-stimulus time histograms for each neuron. There, the approach used trial-averaged histograms making *within-trial* comparisons on a *seconds* timescale. Here, the spike count per trial during baseline window and response windows are used, which allows us to compare across epochs, making *between-trial* comparisons of a 1 second window across the session on a *minutes* timescale. We performed hierarchical clustering on the spike count per trial over all neurons and identified 13 distinct clusters (Figure 7b, c). We used linear regression to identify if a trend was present during the ITI in either the reward and punishment protocol. We found a significant change across epochs in half of the clusters we identified.

**Fig. 7.**
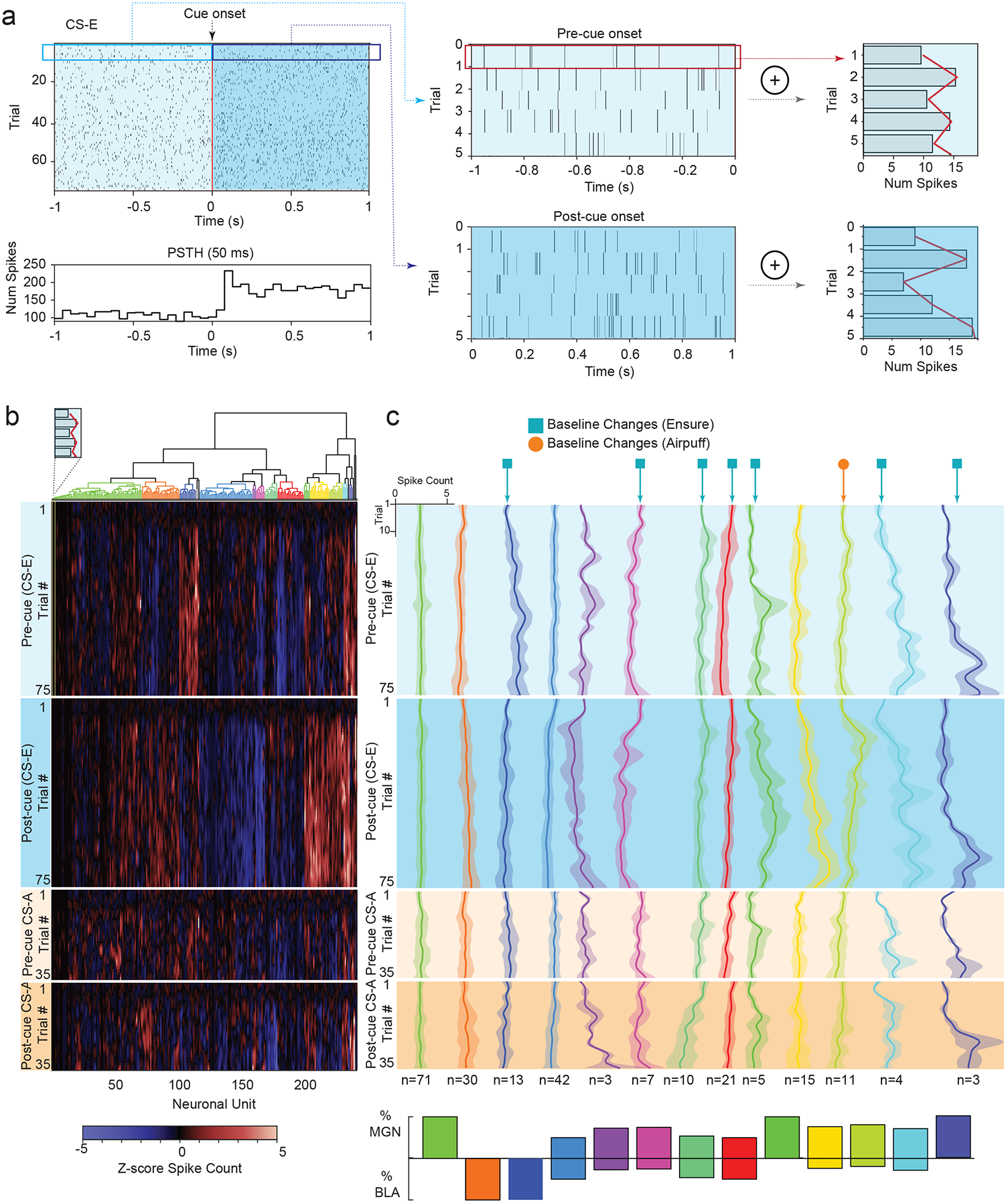
Hierarchical Clustering Across Trials Highlight MGN Neurons that Exhibit Tonic Changes across Trials during Baseline ITI. (a) Schematic plot highlights baseline and response windows used to compute spike counts per trial. Leftmost diagrams depict raster (top) and typical trial-averaged PSTH (bottom). Detail of spike raster for first five trials during baseline (top middle) and response windows (bottom middle) and the summed count for each trial of the baseline (top right) and response (bottom right). (b) Hierarchical clustering on normalized spike counts, summed across time dimension during specific time periods. Dendrogram along top shows resulting clusters; heatmap shows 2 regions for each CS protocol, 4 conditional cases in all: 75 trials for pre-stimulus period (−1 second to cue onset) for CS-E, 75 trials for post-stimulus period (cue onset to +1 second) for CS-E; and the pre-cue baseline and post-cue response for the 35 trials for CS-A. Neuronal units run along x-axis; conditional cases showing trial number make up the y-axis. Dark bands at the upper portion of each conditional case are the result of z-score normalization using the 10-trial habituation period as baseline. (c) Mean z-score response by trial number for the four conditions: reward conditioning shown in blue shading, punishment conditioning shown with orange shading with lighter shade representing pre-cue onset, and darker representing post cue-onset. Inset axes show scale of z-score responses by trial. Blue square identifies clusters with tonic changes during the pre-cue baseline period before CS-E trials, while orange circle marker identifies clusters with tonic change during baseline preceding CS-A. Clusters containing a single neuron across conditions are not shown. Tonic change defined by a non-zero slope of regression line fit to mean response of all units within cluster. Inset bar plot shows percentage of MGN or BLA cells contained within each cluster.

### Summary of Results

In brief, we have seen that while MGN cells are significantly excited to either the reward or punishment stimulus compared to other *a priori* categories, and that BLA cells have no significant difference between categories (Figure 3d). We compared two *a priori* categories (E+A+, E-A-) between their MGN subpopulations and found no significant differences in either case; however, the BLA does show a difference between In-Network and Out-of-Network subpopulations for E-A-(Figure 3e). We adopted an unsupervised clustering approach to explore responses to CS-E and CS-A and found that a variety of response profiles were represented in the MGN but fewer of these response profiles were represented in the BLA (Figure 4e, f). We used the same clustering approach on data that explored CS-E and CS-A sorted by correct or incorrect behavioral responses into Hit, Miss, Correct Rejection, False Alarm categories and found a similar trend; the MGN represented all clusters while the BLA contained fewer than half the number of clusters (Figure 5e, f). Moreover, the neural trajectories for these four categories presented different dynamics in the two regions. In the MGN, Correct Rejection was the longest trajectory and was furthest from Hit; however, in the BLA, there was no significant difference between trajectory lengths and the greatest distance was instead between Hit and False Alarm (Figure 5a, b). Finally, we also uncovered evidence of slower, tonic changes during baseline periods using a novel, albeit elementary, method to dissect trial-by-trial changes (Figure 7).

In conclusion, we have identified strong evidence that robust signals sent from the MGN to the BLA largely carry valence-independent signals that refer to the “absolute value” or salience of the stimulus, while the lower amplitude signals found in the BLA reveal some valence coding populations. In summary, there is filtering that occurs at the level of transmission between the MGN and BLA that enables this transformation, which we speculate may include axo-axonal connections, heavy local inhibition and summation of other inputs, dendritic nonlinearities or shunting. Further, we show that in both the BLA and MGN, we see tonic changes in addition to changes in phasic responses to discrete cues that may reflect changes in internal state or contextual modulation.

passing through the MGN to the BLA appears to undergo a subtle filtering based principally on salience or arousal rather than valence.

## Discussion

There is a rich history in studying Pavlovian fear conditioning in the amygdala to better understand the cellular mechanisms underlying learning. Due to the simple experimental design, robust results, and repeatability, this paradigm has been widely studied at the anatomical (LeDoux, 1986; Yaniv et al., 2001) and circuit levels (Romanski and LeDoux, 1992), and in reference to synaptic pathways (Duvarci and Pare, 2014; Maren, 2005). There are two ways that sensory information enters the amygdala: from thalamic input or via cortical input pathways. Cortical input is generally considered to be slower and more precise, whereas the thalamus represents the rapid relay of sensory information (Maren and Quirk, 2004). For associative conditioning to occur, there must be NMDAR-dependent long-term potentiation in the amygdala (Clem and Huganir, 2010; Janak and Tye, 2015; McKernan and Shinnick-Gallagher, 1997; Rodrigues et al., 2001; Rogan et al., 1997; Rumpel et al., 2005; Tye et al., 2008). However, given that these processes also occur in the thalamus, theoretically the thalamus could be synthesizing and filtering this information before passing it to the amygdala, and acting as more than a simple relay (Jones, 1991; Lee et al., 2010; Murray Sherman, 2001; Sherman, 2007; Weinberger, 2011).

We have uncovered strong evidence in support of the notion that the thalamus is indeed more than a simply relay system as suggested by others. First, the MGN is filtering salient information to the BLA and transforming other valence-enriched signals in the MGN. Therefore, we postulate that the thalamoamygdala circuit follows a model described by the Two-Factor Theory of Emotion (Aron et al., 2005; Schachter and Singer, 1962; Tye, 2018), where recognition of salience in the thalamus is followed by assignment of valence downstream in the amygdala. Second, the MGN and the BLA process expected outcomes differently, as evidenced by correct vs. incorrect behavioral response. Dynamic changes in firing rates accompany anticipatory punishment in the MGN whereas in the BLA, these greater dynamics instead follow unexpected punishments. Third, we have implemented a simple method to identify tonic changes that occur during inter-trial intervals, dynamics which previously have not been studied in detail.

### MGN filtering salient information directly to BLA but routing valence-assigned signals via indirect pathways to BLA

In this study, an optogenetic technique known as phototagging allowed us to identify four populations: (1) cells that project from the MGN to the BLA (MGN→BLA), (2) non-specific MGN cells, (3) cells in the BLA that fire in response to light-evoked activation of the MGN within a short latency (In-Network), and (4) those that do not (Out-of-Network BLA). First, we found that the BLA has proportionally more In-Network neurons that respond to both CS and US than the Out-of-Network populations; however, there is no proportional difference for MGN→BLA and non-specific cells in the MGN (Figure 3c). Second, we defined explicit, *a-priori* response categories and observed how the neurons in each brain region responded. We found evidence that all MGN cells overwhelmingly exhibited a valence-independent excitatory response profile. By contrast, the BLA did not show a statistically significant difference in proportion across the different categories (Figure 3d). Within the preferred category, we did not find notable differences in the proportion of MGN→BLA or non-specific MGN cells. However, the BLA did show significant differences between In-Network and Out-of-Network populations across the largest categories (Figure 3e). While it is hardly surprising that the MGN cells respond to salience, it is interesting to note the MGN Phototagged cells exhibited less functional diversity than non-phototagged cells. Certain responses that differentiate positive and negative valence actually disappear in the MGN→BLA population (E+A-, E-, and A-) while others are richly represented. Somehow, the large amplitude, sharp transient responses in MGN cells are not present in the BLA – either through presynaptic modulation, dendritic filtering, or local inhibition within the BLA, these robust signals are either muted or absent in the BLA (Figure 3e). Why they exist in MGN→BLA neurons but not in BLA neurons receiving input from MGN→BLA neurons is unclear. Perhaps they send collaterals to the cortex or other regions where the sharp amplitude signals are propagated, while axo-axonal, dendritic or somatic inhibition may contribute to the most robust signals being filtered out before arriving within the BLA.

Another interesting note is that the individual *a priori* categories are represented in MGN→BLA and In-Network BLA subpopulations to different degrees. This is a surprising result, as these two populations directly share information (Figure 3e). We see evidence of the A-category in In-Network BLA yet the same category is absent entirely in the MGN→BLA subpopulation. One could interpret this by noting the importance of the polysynaptic connections that make up the In-Network population; information is propagating into the BLA but not, apparently, via direct MGN→BLA connections. The thalamo-cortico-amygdala pathway, which plays a strong role in memory and fear learning (Ferrara et al., 2017), could be collateralizing signals. Of the two signaling stream pathways in the thalamocortical system, the higher order pathway does appear to be collateralized; we speculate here that by sending the signal to multiple brain regions, one region is connecting with interneurons that cancels out the direct signals coming from thalamus (Wolff, 2021), resulting in different response profiles in evidence in the BLA. Alternatively, input sent along direct projections into the BLA could undergo dendritic filtering, which exists to intentionally dampen strong signals coming into the amygdala and is likely to preserve signals with highest salience.

Our general interpretation of these data is that the BLA represents a diverse response that incorporates both arousal and valence while the MGN primarily represents arousal. This result follows one of two models highlighted in a recent review article (Tye, 2018), specifically, the Two-Factor Theory of Emotion. Within the framework of valence processing, the Two-Factory Theory outlines the following order of operations for two distinct neural processes: first, the brain must recognize the salience of a stimulus as the overall magnitude of intensity of the response (|*n*|) before assigning a positive or negative valence to the stimulus (+*n* or -*n*). Here, we see indication of pure salience occurring upstream in the MGN, where information is largely independent of valence, and valence is assigned downstream in the BLA. To our knowledge, this is the first time this model has been validated through anatomical structure and function. Moreover, although diverse response categories are present in the MGN (Figures 3e, 4e-h, 5e-h), there is not a one-to-one match across MGN→BLA and In-network populations indicating that the MGN is routing information via alternate, indirect pathways that ultimately terminate in the BLA.

### MGN and BLA respond differently to expected and unexpected outcomes

We investigated the response of the MGN and the BLA to account for different representations when behavior matched or contrasted with the expected outcome of the tone (hit, miss, correct rejection, false alarm) by plotting the neural trajectories in reduced dimensional space. In the MGN, there was a significantly longer trajectory associated with Correct Rejection and significantly greater distances between Hit and Correct Rejection trajectories than most other pairings. This indicates more dynamics for these two conditions and fewer dynamic differences during Miss or False Alarm (Figure 5a). By contrast, in the BLA, although trajectories are of similar length, their relative distances have shifted such that Hit and False alarm are further apart whereas Miss and Correct Rejection are less dynamic. Why would the MGN appear to represent Hit and Correct Rejection with different dynamics compared to the other responses, while the BLA minimizes the role of Correct Rejection and instead emphasize False Alarm with Hit?

One interpretation of these findings is that error signals are more pronounced in the BLA, while expectation of punishment is amplified in the MGN. The MGN projects to multiple layers within the cortex (Lee, 2015) and the thalamocortical pathway was found to be the principal fear memory pathway of an intact brain (Boatman et al., 2006). Thus, it follows that anticipatory knowledge of a coming punishment is amplified in the MGN as this signal is likely sent to cortical regions. However, since False Alarm is paired with an unexpected punishment, an incorrectly identified signals would receive greater emphasis in the BLA. In contrast to the classic reward prediction error signals seen in some VTA dopamine neurons(Schultz et al., 1997), a subset of BLA neurons have actually been shown to exhibit a phasic excitation in response to unexpected reward omission that is correlated to frustrative nonreward. In this study, frustration was operationally defined as an increase in vigor or intensity of responding following an unexpected reward omission in an operant conditioning task (Tye et al., 2010).

### Evidence of both Phasic and Tonic Changes

Strong phasic responses within an experimental response window are the basis for investigating neural responses to stimuli. Frequently, researchers average across multiple trials to form a detailed response profile in time. Although working with trial-averaged responses has a long and successful practice, averaging obscures slow-scale changes that could emerge during the learning process. In this paper, we introduced a novel method for investigating slow-scale, tonic changes that accompany learning. Instead of averaging across trials, we drill down into the trial-by-trial changes by clustering spike counts. We identified subsets of neurons that experience slow but observable changes to their firing rates across the trials. More surprisingly, we uncovered changes during baseline periods, which are traditionally assumed to be static. Tonic firing more often consists of a steady firing at a specific frequency so changes to this background state that accompany learning are an unexplored area. It is our opinion that investigating trial-by-trial changes is a potentially rich area of study, which we intend to study further with increased specificity and granularity.

### Summary and Future Directions

In the current study, we chose to focus on direct projections between the MGN and BLA, which have a specific response profile, namely excitatory long-range responses that propagate downstream to other brain regions. However, the thalamus, and specifically the MGN, has a diverse representation of other types of cells such as interneurons that might have firing rates orders of magnitude higher than projectors. Another important future direction will be to observe the activity of MGN interneurons or other locally-projecting neurons which could have distinct activity patterns involved in filtering and processing of information distinct from MGN→BLA cells.

We have demonstrated that the role of the MGN in relaying sensory information is not a static one. It appears to aid the learning process by filtering incoming sensory information to route saliency information directly to the BLA and other information, such as error signaling, indirectly to the BLA. While additional work will be necessary to evaluate the validity of our findings, our work supports the two factor theory of emotion with MGN routing arousal information to the BLA, and suggests the exploration of computations in the MGN that shape learning, as opposed to treating MGN as a sensory relay.

## Acknowledgments

KT is an HHMI Investigator and the Wylie Vale Professor at the Salk Institute for Biological Studies, and this work was supported by funding from Salk, HHMI, Clayton Foundation, Kavli Foundation, Dolby Family Fund, R01-MH115920 (NIMH), R37-MH102441 (NIMH), and Pioneer Award DP1-AT009925 (NCCIH). JMO was supported by a Brain Initiative F32 from NIMH (F32 MH115446-01).

## AUTHOR CONTRIBUTIONS

CL and KT conceptualized the project and designed the experiments. CL performed preprocessing components of data analysis for the behavioral and in vivo electrophysiology data. LK performed statistical analyses and hierarchical clustering analyses. LK and KB analyzed neural trajectories. LK conceptualized and implemented hierarchical clustering of spike count data and implemented state-space analysis. GM, GG, MJ, YF, AB, and CL contributed to data collection. JO, EN, CW, RW and EK contributed technical expertise in establishing hardware or software used for data collection. LK, CL, and KT wrote the manuscript, CL, GM, and LK plotted the figures, all authors contributed to revising the manuscript. KT supervised all stages of the work.

These authors contributed equally: Chris Leppla, Laurel Keyes

## Figure Legends

**Supplemental Fig. 1.**
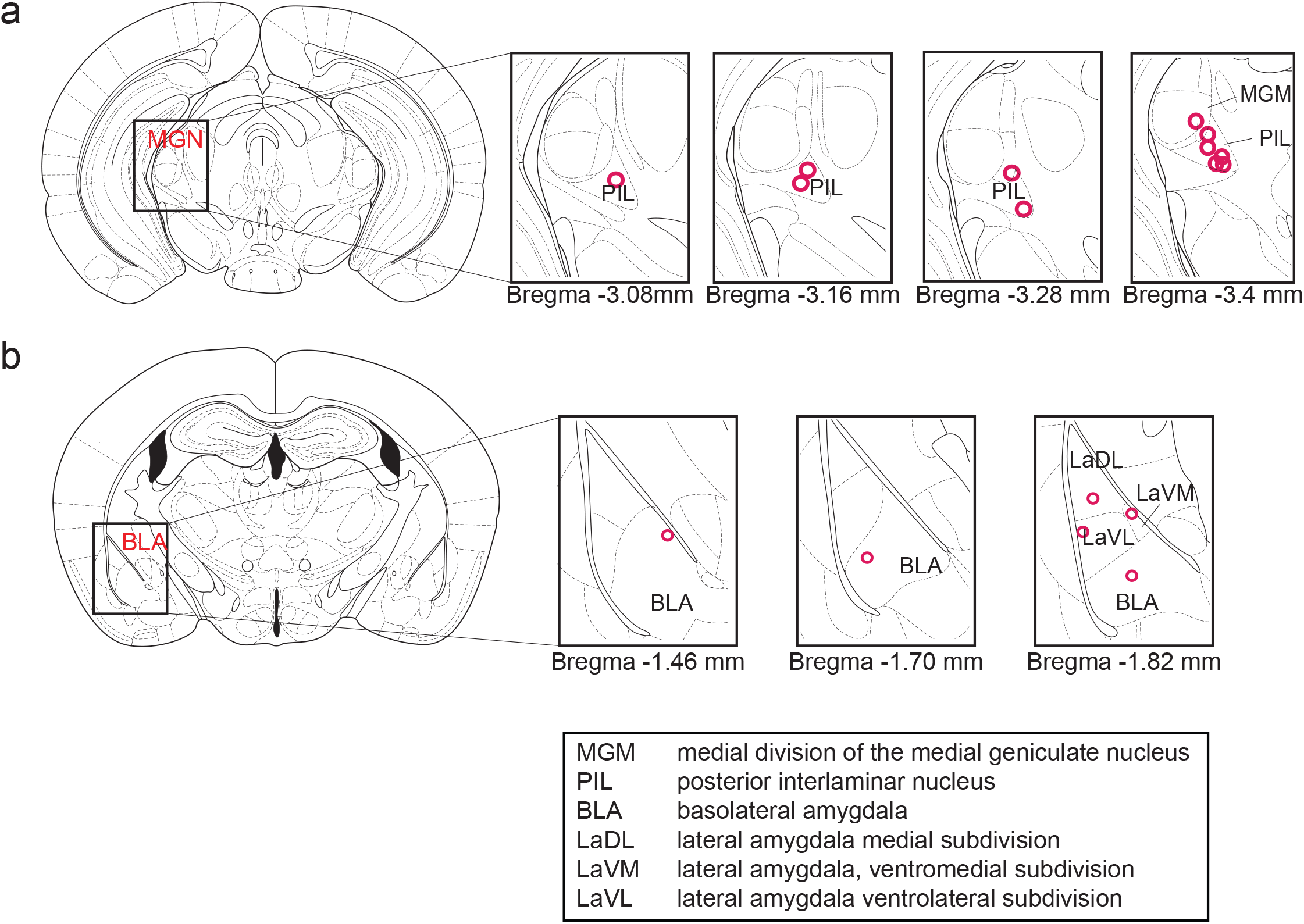
Viable Electrode Placements in MGN and BLA for electrophysiology recordings. (a, b) Electrolytic lesion sites indicate recording location in MGN (top) and BLA (bottom). Experimental subjects with optrode placements outside of MGN were excluded from analysis. Subjects with correct electrode placements in MGN did not all have viable electrode placements in BLA. Electrode placements outside BLA are not shown, as data from these channels was excluded from analysis.

**Supplemental Fig. 2.**
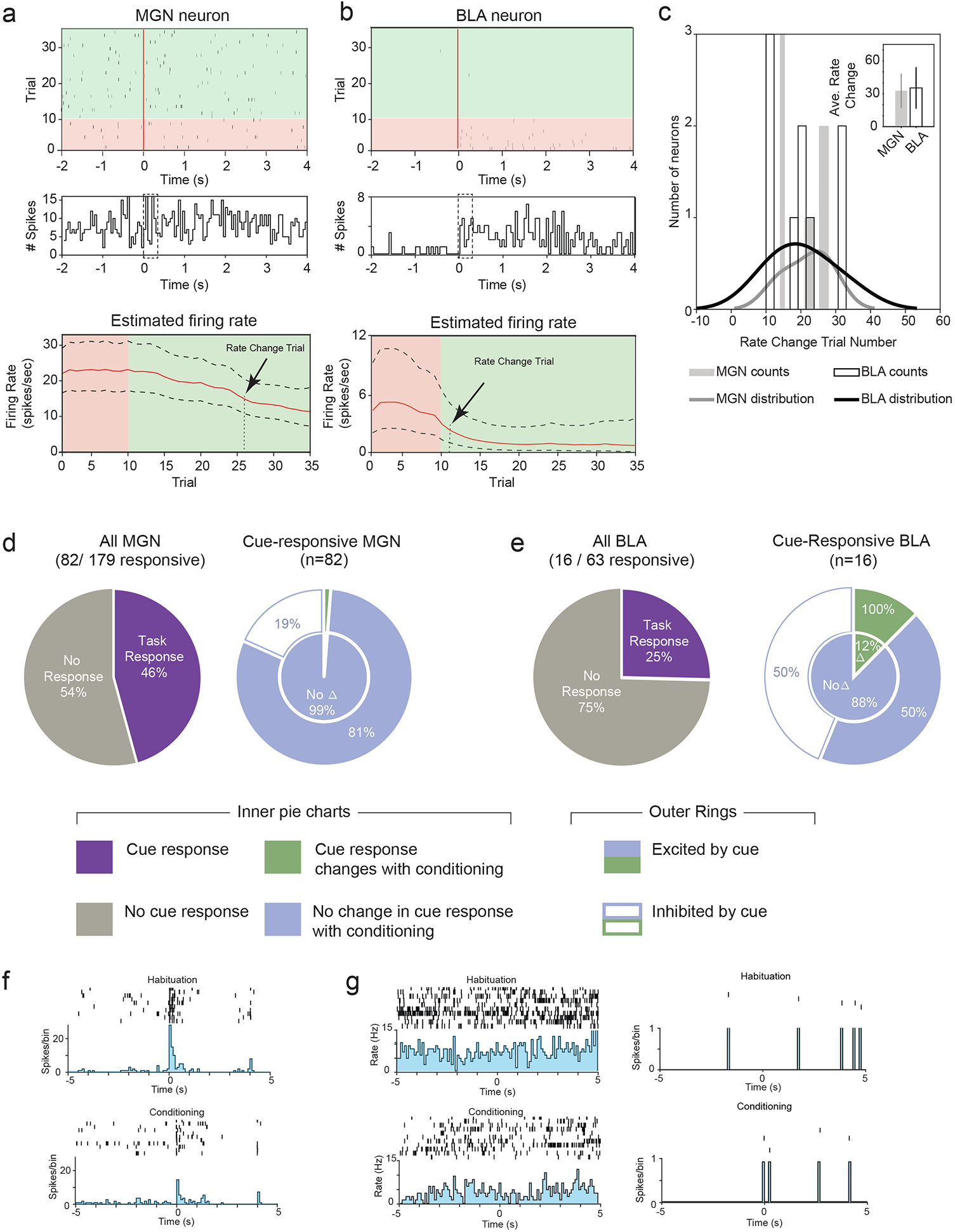
MGN and BLA Neurons Exhibit Similar Rate Change Trial During Discrimination Learning. (a, b) Representative raster and PSTH (50 ms bins) for the period −2 s before cue to +4 s after cue for (a) MGN and (b) BLA neuron; baseline early trials (shaded red) and conditioning (shaded green) periods, tone onset shown by red line. Dotted box in PSTH represents the 150 ms interval over which state space analysis was computed. Lower plots show probabilistic estimate of the trial at which neuron undergoes a rate change using state-space analysis; black arrow indicates rate change trial number. (c) Representative example of randomly selected trial matched pairs of rate change trial number for MGN and BLA neurons for CS-A. The distributions do not exhibit separation in the rate change trial (Kolmogorov–Smirnov test inconclusive for 25 randomly selected matched trials); thus, our findings are inconclusive as to whether one neuronal population learns before the other. (d) Cue-responsive MGN subpopulations; 46% of cells respond to task. Of these,1% (n=1) changed with conditioning. (e) Cue-responsive BLA subpopulations; 25% responded to task. Of these 12% (n=2) changed with conditioning. (f) Spike raster and PSTH for single MGN cell that changed with conditioning. This cell was initially excited to cue and weakened with conditioning. (g) Spike raster and PSTH for both BLA cells that changed with conditioning. Both cells were initially excited to cue and weakened with conditioning.

## Supplemental Metshods

### MGN and BLA Neurons Exhibit Similar Encoding Rates During Discrimination Learning and Few Neurons Exhibit Conditioning during Punishment Learning

We applied a state-space analysis to all neurons in each brain region in order to characterize the neural dynamics in the MGN and the BLA (Supplemental Figure 2a, b). Such models allow accurate estimation of instantaneous firing rate as well as the ability to identify the trial at which the firing rate changes within a statistical framework; the latter “rate-change trial” represents the point at which a neuron encodes the learned association. We hypothesized that the MGN would encode this information before the BLA but found no significant difference between their rate change trial distributions (Kolmogorov–Smirnov test, matched pairs, n.s., Supplemental Figure 2d). In other words, the MGN does not appear to learn an association before passing on information downstream.

To characterize whether the neural response changed over the course of the discrimination session, early trials and late trials of discrimination session data were compared (Supplemental Figure 2d). Due to limitations in the experimental paradigm, a habituation phase was not expressly included in the trial structure. To interrogate whether neurons changed with conditioning, we denote trials 1-10 of the punishment cue to act as the habituation phase during discrimination learning, while trials 11-35 form the conditioning phase. There were proportionally more cue-responsive neurons in the MGN than in the BLA (MGN: 46%, 82/179, N = 12; BLA: 25%, 16/63, N = 6). Of these cue-responsive neurons, proportionally fewer MGN neurons experienced a change with conditioning than BLA neurons (MGN: 1%, 1/82, N = 12; BLA: 12%, 2/16, N=6). The sole MGN neuron that underwent a change in conditioning was excited to the cue and the mean z-score response weakened with conditioning (Supplemental Figure 2f). The two BLA neurons that changed with conditioning were both excited to the cue and weakened the magnitude response with conditioning (Supplemental Figure 2g).

### Calculation of firing rate and rate change trial

Firing rate was modeled using the state-space approach described in Smith et al. (2010) and was modified as described in (Allsop et al., 2018) to find the rate change trial. The MATLAB code for computing state-space analysis of neural firing may be downloaded from http://annecsmith.net/firingrates.html. To determine whether a cell changed with conditioning over the course of the task, the first 10 trials of punishment data were assigned as the habituation phase and the last 10 trials as the conditioning phase. An ANOVA across four time-windows (two baseline and two response) was computed to see if there was a difference between the average firing rate in each window. We then compared the two experimental windows in post-hoc comparison tests. If there was a statistically significant difference between the firing rate across both experimental response windows, that unit was identified as changing with conditioning (Task Δ).

## REFERENCES

Allsop SA, Wichmann R, Mills F, Burgos-Robles A, Chang C-J, Felix-Ortiz AC, Vienne A, Beyeler A, Izadmehr EM, Glober G, Cum MI, Stergiadou J, Anandalingam KK, Farris K, Namburi P, Leppla CA, Weddington JC, Nieh EH, Smith AC, Ba D, Brown EN, Tye KM. 2018. Corticoamygdala Transfer of Socially Derived Information Gates Observational Learning. Cell 173:1329-1342.e18. doi:10.1016/j.cell.2018.04.004

Antunes R, Moita MA. 2010. Discriminative Auditory Fear Learning Requires Both Tuned and Nontuned Auditory Pathways to the Amygdala. J Neurosci 30:9782–9787. doi:10.1523/JNEUROSCI.1037-10.2010

Aron A, Fisher H, Mashek DJ, Strong G, Li H, Brown LL. 2005. Reward, Motivation, and Emotion Systems Associated With Early-Stage Intense Romantic Love. J Neurophysiol 94:327–337. doi:10.1152/jn.00838.2004

Beyeler A, Chang C-J, Silvestre M, Lévêque C, Namburi P, Wildes CP, Tye KM. 2018. Organization of Valence-Encoding and Projection-Defined Neurons in the Basolateral Amygdala. Cell Reports 22:905–918. doi:10.1016/j.celrep.2017.12.097

Beyeler A, Namburi P, Glober GF, Simonnet C, Calhoon GG, Conyers GF, Luck R, Wildes CP, Tye KM. 2016. Divergent Routing of Positive and Negative Information from the Amygdala during Memory Retrieval. Neuron 90:348–361. doi:10.1016/j.neuron.2016.03.004

Bordi F, LeDoux JE. 1994. Response properties of single units in areas of rat auditory thalamus that project to the amygdala. II. Cells receiving convergent auditory and somatosensory inputs and cells antidromically activated by amygdala stimulation. Exp Brain Res 98:275–286. doi:10.1007/BF00228415

Boyden ES, Zhang F, Bamberg E, Nagel G, Deisseroth K. 2005. Millisecond-timescale, genetically targeted optical control of neural activity. Nat Neurosci 8:1263–1268. doi:10.1038/nn1525

Burgos-Robles A, Kimchi EY, Izadmehr EM, Porzenheim MJ, Ramos-Guasp WA, Nieh EH, Felix-Ortiz AC, Namburi P, Leppla CA, Presbrey KN, Anandalingam KK, Pagan-Rivera PA, Anahtar M, Beyeler A, Tye KM. 2017. Amygdala inputs to prefrontal cortex guide behavior amid conflicting cues of reward and punishment. Nat Neurosci 20:824–835. doi:10.1038/nn.4553

Chaudhri N, Sahuque LL, Janak PH. 2008. Context-Induced Relapse of Conditioned Behavioral Responding to Ethanol Cues in Rats. Biological Psychiatry 64:203–210. doi:10.1016/j.biopsych.2008.03.007

Chaudhri N, Sahuque LL, Schairer WW, Janak PH. 2009. Separable roles of the nucleus accumbens core and shell in context-and cue-induced alcohol-seeking. Neuropsychopharmacology 35:783–791.

Clem RL, Huganir RL. 2010. Calcium-Permeable AMPA Receptor Dynamics Mediate Fear Memory Erasure. Science 330:1108–1112. doi:10.1126/science.1195298

Cruikshank SJ, Edeline JM, Weinberger NM. 1992. Stimulation at a site of auditory-somatosensory convergence in the medial geniculate nucleus is an effective unconditioned stimulus for fear conditioning. Behav Neurosci 106:471–483. doi:10.1037//0735-7044.106.3.471

Duvarci S, Pare D. 2014. Amygdala Microcircuits Controlling Learned Fear. Neuron 82:966–980. doi:10.1016/j.neuron.2014.04.042

Edeline J-M, Neuenschwander-El Massioui N, Dutrieux G. 1990. Discriminative long-term retention of rapidly induced multiunit changes in the hippocampus, medial geniculate and auditory cortex. Behavioural Brain Research 39:145–155. doi:10.1016/0166-4328(90)90101-J

Fenno L, Yizhar O, Deisseroth K. 2011. The Development and Application of Optogenetics. Annual Review of Neuroscience 34:389–412. doi:10.1146/annurev-neuro-061010-113817

Ferrara NC. 2015. Neural Mechanisms Supporting Differential Auditory Fear Conditioning. Theses and Dissertations 1048.

Han J-H, Yiu AP, Cole CJ, Hsiang H-L, Neve RL, Josselyn SA. 2008. Increasing CREB in the auditory thalamus enhances memory and generalization of auditory conditioned fear. Learning & Memory 15:443–453. doi:10.1101/lm.993608

Janak PH, Tye KM. 2015. From circuits to behaviour in the amygdala. Nature 517:284–292. doi:10.1038/nature14188

Jones EG. 1991. Chapter 3 The anatomy of sensory relay functions in the thalamus In: Holstege G, Editor. Progress in Brain Research, Role of The Forebrain in Sensation and Behavior. Elsevier. pp. 29–52. doi:10.1016/S0079-6123(08)63046-0

LeDoux JE. 2000. Emotion Circuits in the Brain. Annual Review of Neuroscience 23:155–184. doi:10.1146/annurev.neuro.23.1.155

LeDoux JE. 1986. Sensory systems and emotion: A model of affective processing. Integrative Psychiatry 4:237–243.

Lee S-C, Cruikshank SJ, Connors BW. 2010. Electrical and chemical synapses between relay neurons in developing thalamus. The Journal of Physiology 588:2403–2415. doi:10.1113/jphysiol.2010.187096

Lima SQ, Hromádka T, Znamenskiy P, Zador AM. 2009. PINP: A New Method of Tagging Neuronal Populations for Identification during In Vivo Electrophysiological Recording. PLoS ONE 4:1–10. doi:10.1371/journal.pone.0006099

Maren S. 2005. Synaptic mechanisms of associative memory in the amygdala. Neuron 47:783–786. doi:10.1016/j.neuron.2005.08.009

Maren S, Quirk GJ. 2004. Neuronal signalling of fear memory. Nat Rev Neurosci 5:844–852. doi:10.1038/nrn1535

McKernan MG, Shinnick-Gallagher P. 1997. Fear conditioning induces a lasting potentiation of synaptic currents in vitro. Nature 390:607–611. doi:10.1038/37605

Murray Sherman S. 2001. Chapter 4 Thalamic relay functionsProgress in Brain Research, Vision: From Neurons to Cognition. Elsevier. pp. 51–69. doi:10.1016/S0079-6123(01)34005-0

Nabavi S, Fox R, Proulx CD, Lin JY, Tsien RY, Malinow R. 2014. Engineering a memory with LTD and LTP. Nature 511:348–352. doi:10.1038/nature13294

Namburi P, Beyeler A, Yorozu S, Calhoon GG, Halbert SA, Wichmann R, Holden SS, Mertens KL, Anahtar M, Felix-Ortiz AC, Wickersham IR, Gray JM, Tye KM. 2015. A circuit mechanism for differentiating positive and negative associations. Nature 520:675–678. doi:10.1038/nature14366

Nieh EH, Matthews GA, Allsop SA, Presbrey KN, Leppla CA, Wichmann R, Neve R, Wildes CP, Tye KM. 2015. Decoding neural circuits that control compulsive sucrose seeking. Cell 160:528–541. doi:10.1016/j.cell.2015.01.003

Nishijo H, Ono T, Nishino H. 1988. Single neuron responses in amygdala of alert monkey during complex sensory stimulation with affective significance. J Neurosci 8:3570–3583. doi:10.1523/JNEUROSCI.08-10-03570.1988

Orsini CA, Maren S. 2012. Neural and cellular mechanisms of fear and extinction memory formation. Neuroscience & Biobehavioral Reviews, Memory Formation 36:1773–1802. doi:10.1016/j.neubiorev.2011.12.014

Parsons RG, Riedner BA, Gafford GM, Helmstetter FJ. 2006. The formation of auditory fear memory requires the synthesis of protein and mRNA in the auditory thalamus. Neuroscience 141:1163–1170. doi:10.1016/j.neuroscience.2006.04.078

Rodrigues SM, Schafe GE, LeDoux JE. 2001. Intra-Amygdala Blockade of the NR2B Subunit of the NMDA Receptor Disrupts the Acquisition But Not the Expression of Fear Conditioning. The Journal of Neuroscience 21:6889–6896.

Rogan MT, Stäubli UV, LeDoux JE. 1997. Fear conditioning induces associative long-term potentiation in the amygdala. Nature 390:604–607. doi:10.1038/37601

Romanski LM, LeDoux JE. 1992. Equipotentiality of thalamo-amygdala and thalamo-cortico-amygdala circuits in auditory fear conditioning. J Neurosci 12:4501–4509.

Rumpel S, LeDoux J, Zador A, Malinow R. 2005. Postsynaptic Receptor Trafficking Underlying a Form of Associative Learning. Science 308:83–88. doi:10.1126/science.1103944

Rushton W a. H. 1927. The effect upon the threshold for nervous excitation of the length of nerve exposed, and the angle between current and nerve. The Journal of Physiology 63:357–377. doi:10.1113/jphysiol.1927.sp002409

Schachter S, Singer JE. 1962. Cognitive, social, and physiological determinants of emotional state. Psychol Rev 69:379–399.

Schultz W, Dayan P, Montague PR. 1997. A Neural Substrate of Prediction and Reward. Science 275:1593–1599. doi:10.1126/science.275.5306.1593

Sciascia JM, Reese RM, Janak PH, Chaudhri N. 2015. Alcohol-Seeking Triggered by Discrete Pavlovian Cues is Invigorated by Alcohol Contexts and Mediated by Glutamate Signaling in the Basolateral Amygdala. Neuropsychopharmacol 40:2801–2812. doi:10.1038/npp.2015.130

Senn V, Wolff SBE, Herry C, Grenier F, Ehrlich I, Gründemann J, Fadok JP, Müller C, Letzkus JJ, Lüthi A. 2014. Long-range connectivity defines behavioral specificity of amygdala neurons. Neuron 81:428–437. doi:10.1016/j.neuron.2013.11.006

Sherman SM. 2007. The thalamus is more than just a relay. Current Opinion in Neurobiology, Sensory systems 17:417–422. doi:10.1016/j.conb.2007.07.003

Tye KM. 2018. Neural Circuit Motifs in Valence Processing. Neuron 100:436–452. doi:10.1016/j.neuron.2018.10.001

Tye KM, Cone JJ, Schairer WW, Janak PH. 2010. Amygdala Neural Encoding of the Absence of Reward during Extinction. The Journal of Neuroscience 30:116–125. doi:10.1523/JNEUROSCI.4240-09.2010

Tye KM, Deisseroth K. 2012. Optogenetic investigation of neural circuits underlying brain disease in animal models. Nat Rev Neurosci 13:251–266. doi:10.1038/nrn3171

Tye KM, Stuber GD, De Ridder B, Bonci A, Janak PH. 2008. Rapid strengthening of thalamo-amygdala synapses mediates cue–reward learning. Nature 453:1253–1257.

Valyear MD, Glovaci I, Zaari A, Lahlou S, Trujillo-Pisanty I, Andrew Chapman C, Chaudhri N. 2020. Dissociable mesolimbic dopamine circuits control responding triggered by alcohol-predictive discrete cues and contexts. Nature communications 11:1–14.

Weinberger NM. 2011. The medial geniculate, not the amygdala, as the root of auditory fear conditioning. Hearing Research 274:61–74. doi:10.1016/j.heares.2010.03.093

Wepsic JG. 1966. Multimodal sensory activation of cells in the magnocellular medial geniculate nucleus. Experimental Neurology 15:299–318. doi:10.1016/0014-4886(66)90053-7

Yaniv D, Schafe GE, LeDoux JE, Richter-Levin G. 2001. A gradient of plasticity in the amygdala revealed by cortical and subcortical stimulation, in vivo. Neuroscience 106:613–620. doi:10.1016/s0306-4522(01)00312-8

